# Feature detecting columnar neurons mediate object tracking saccades in *Drosophila*

**DOI:** 10.1101/2022.09.21.508959

**Authors:** Giovanni Frighetto, Mark A. Frye

## Abstract

Tracking visual objects while stabilizing the visual background is complicated by the different computational requirements for object tracking and motion vision. In fruit fly, directionally selective motion detectors T4 and T5 cells supply wide-field neurons of the lobula plate, which control smooth gaze stabilization behavior. Here, we hypothesized that an anatomically parallel pathway supplied by T3, which encodes small moving objects and innervates the lobula, drives body saccades toward objects. We combined physiological and behavioral experiments to show that T3 neurons respond omnidirectionally to contrast changes induced by the visual stimuli that elicit tracking saccades, and silencing T3 reduced the frequency of tracking saccades. By contrast, optogenetic manipulation of T3 increased the number of tracking saccades. Our results represent the first evidence that parallel motion detection and feature detection pathways coordinate smooth gaze stabilization and saccadic object tracking behavior during flight.

## Introduction

Discriminating and tracking a moving visual object against a cluttered and moving visual background represents a complex task that nevertheless is common across taxa. Flies have shown a stunning ability to actively orient toward and fixate foreground stimuli whose detection relies on very few parameters and is remarkably robust to perturbation (Egelhaaf, 1985; Reichardt et al., 1983; Theobald et al., 2008). Object orientation behavior becomes extremely challenging during locomotion, when the visual panorama moves across the retina generating complex patterns of optic flow that can move with or against the direction of the pursued object. *Drosophila* has been a productive model for understanding how patterns of wide-field optic flow are decoded by an array of small-field local motion detectors (Borst, 2014; Mauss and Borst, 2020) and how local motion direction is computed (Borst et al., 2020b; Groschner et al., 2022; Gruntman et al., 2019). Local motion detectors segregate into parallel ON and OFF luminance selective cells T4 and T5, respectively (Joesch et al., 2010; Strother et al., 2017). These two types of columnar neurons have four subtypes, each tuned to a singular cardinal direction, which each innervate one of four direction-selective layers of the lobula plate, the fourth neuropil of the optic lobe (Fisher et al., 2015; Maisak et al., 2013). Second-order interneurons of the lobula plate pool inputs from T4/T5 cells to assemble complex spatial filters for patterns of optic flow, and then project to pre-motor descending pathways to coordinate syn-directional head and wing steering movements that stabilize gaze against perturbations during locomotion (Busch et al., 2018; Haikala et al., 2013; Heisenberg et al., 1978). However, wide-field optic flow and its corollaries (e.g., looming stimuli) are not the only visual features that flies need to extract from the visual world (Borst et al., 2020a; Cheong et al., 2020).

Identifying foreground objects from a cluttered background requires visual computations that are unlikely to be carried out by the same direction selective system (Aptekar and Frye, 2013; Reichardt et al., 1983). In tethered walking flies, synaptic suppression of T4/T5 cells has been shown to leave transient orientation response toward a flickering bar essentially unaffected, while dramatically reducing the response to a moving bar (Bahl et al., 2013). In a follow-up experiment, flying flies were presented with a textured vertical bar revolving around a circular arena. Flies responded with a distinct initial steering response oriented counter directional to the moving bar, followed by a secondary response in the same direction of bar movement (Keleş et al., 2018). Silencing T4/T5 under these conditions reduced the secondary syn-directional steering response, but left the initial counter-directional orientation phase intact (Keleş et al., 2018). These studies would suggest the existence of a motion-independent sub-system that mediates object orientation, likely detecting spatial contrast information, operating in parallel to the T4/T5 motion detection pathway (Bahl et al., 2015, 2013). One study proposed that a model based solely on T4/T5-like directional motion detectors and small-field integrators such as so-called ‘figure detecting’ neurons (Liang et al., 2012) can account for the smooth steering movements that drive bar fixation behavior by tethered flies, even under the challenge of opposing background motion (Fenk et al., 2014).

By contrast to rigidly tethered flies, animals tethered to a frictionless magnetic pivot and free to steer in the yaw plane execute rapid body saccades to track an object in the form of a rotating vertical edge or bar. By contrast, they execute smooth steering movements to stabilize a revolving wide-field panorama, interspersed with occasional saccades (Mongeau and Frye, 2017). The dynamics of bar tracking saccades are distinct from those triggered by a wide-field motion (Mongeau and Frye, 2017). Therefore, the evidence for or against a T4/T5-independent mechanism for object tracking might be resolved by considering the differential control of smooth optomotor steering and saccadic reorientation.

Here, we support previous lines of evidence that T4/T5 coordinate smooth optomotor responses for wide-field gaze stabilization, but that a parallel neural pathway supplies the control of object tracking saccades. T3 neurons (Keleş et al., 2020; Tanaka and Clark, 2020) arborize within single columns of the medulla and send axons into layers 2 and 3 of the third neuropil of the optic lobe, the lobula (Fischbach and Dittrich, 1989; Takemura et al., 2013). Using calcium imaging, we demonstrate that T3 neurons respond vigorously to the background-matched motion-defined bars that robustly elicit bar tracking saccades (Mongeau et al., 2019; Mongeau and Frye, 2017). In rigidly tethered flies, hyperpolarizing T3 by genetically expressing an inward rectifying potassium channel (Kir2.1) reduces the initial counter-directional orientation response typically deployed for tracking motion-defined bars. In magnetically tethered flies, hyperpolarizing T3 neurons reduced the number of bar tracking saccades, whereas optogenetic activation by CsChrimson increased them. Finally, we posit a role of T3 in triggering bar tracking saccades through an integrate-and-fire model physiologically inspired by the calcium dynamics of T3 neurons and a control model of saccadic bar tracking (Mongeau and Frye, 2017).

## Results

### T3 neurons are well-tuned to encode motion-defined bars

The lobula is mainly innervated by a class of visual projection neurons (VPNs), the lobula columnar (LC) cells, each type of which project together to the central brain forming bundles of type-specific terminals called optic glomeruli (Aptekar et al., 2015; Panser et al., 2016; Wu et al., 2016). In previous work (Keleş et al., 2020), our lab used an intersectional strategy to generate specific split-Gal4 driver lines for two T-shaped neuron types, T3 and T2a, arborising in the medulla and terminating in layer 2 and 3 of the lobula (Fischbach and Dittrich, 1989; Takemura et al., 2013). The cell bodies of T3 and T2a are caudally located in the space between the medulla and lobula plate neuropiles (Fischbach and Dittrich, 1989).

To characterize responses to vertical bars by T3 neurons, we recorded calcium signals under *in vivo* two-photon excitation imaging with an LED visual stimulus (Reiser and Dickinson, 2008) (**Figure 1A**). We imaged from presynaptic terminals in the lobula of flies expressing GCaMP6f in T3 (**Figure 1B**). Flies were presented with bars moving across the right visual field, ipsilateral to the recording site, of varying direction and contrast polarity (**Figure 1C**). T3 showed increase in calcium activity within individual presynaptic terminals, and the ensemble of T3 dendrites innervating layer 9 of the medulla exhibited a robust retinotopic wave of activation as the bar swept across the retina (**Figure 1C, Video 1**). Broadly consistent with prior results (Keleş et al., 2020), T3 neurons were strongly activated by front-to-back and back-to-front motion of either ON (brighter than background) or OFF (darker than background) bars (**Figure 1D**), with a slight preference for OFF transitions.

**Figure 1.**
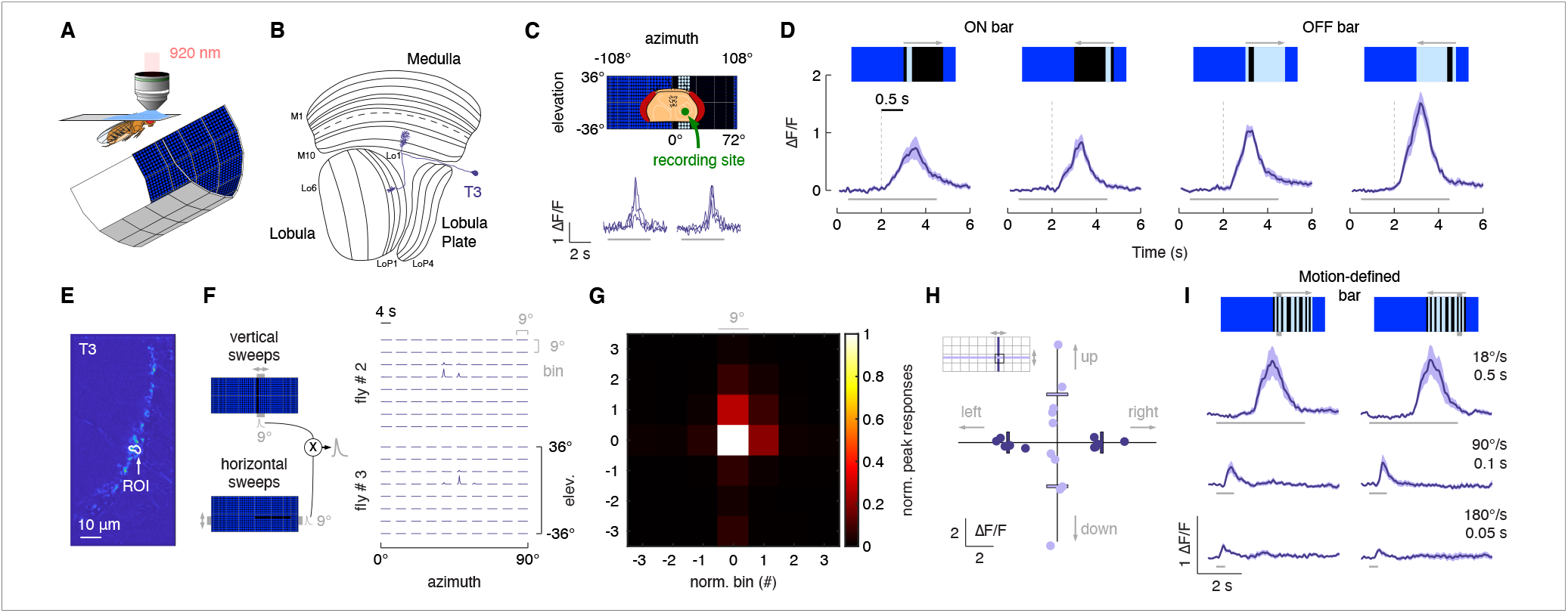
T3s are omnidirectional neurons with small receptive fields and broad temporal sensitivity. (A) Head-fixed fly for two-photon calcium imaging while presented with visual stimuli from a surrounding LED display. (B) Fly optic lobe neuropils (medulla, lobula and lobula plate) with a T3 neuron highlighted in magenta. (C) Top: schematic representation of the fly head and the recording site framed in the center of the LED display. Bottom: calcium imaging responses to a ON solid moving bar at 18° s-1 (left: front-to-back, right: back-to-front) of a T3 neuron from a representative fly (3 repetitions). (D) Average responses (mean ± s.e.m.) to moving ON and OFF solid bars (9° x 72°, width x height) at 18° s-1 in two different directions (front-to-back and back-to-front). Visual stimuli are depicted at the top. Dashed vertical gray lines indicate the coarse onset of the responses. Light gray horizontal bars at the bottom indicate stimulus presentation (n = 11 flies, 3 repetitions per fly). (E) ROI drawn around the presynaptic terminal in the lobula of a T3 neuron expressing GCaMP6f. Image representing the mean activity from the two-photon imaging experiment in a representative fly. (F) Left: representation of the procedure used to compute the RF of T3. Gray shaded region behind the LED display represents a bin (9° x 72°, width x height) within which a single pixel dark bar (2.25° width) is swept in two different directions. The cross product is then obtained by multiplying the calcium responses to vertical and horizontal sweeps. Right: matrix of the multiplied traces in the 10 × 8 bins (horizontal x vertical) in two representative flies. The RF is probed within a window of 90° x 72° (horizontal x vertical). (G) Mean of the normalized peak responses of T3 neurons by spatial location (n = 5 flies). Bin = 0 represents the center of the RF. (H) Directional calcium peak responses to a 2.25° dark bar moving (18° s-1) in the four cardinal directions of individual flies. Bars indicate the mean. (I) Average responses (mean ± s.e.m.) to motion-defined bars (9° x 72°, width x height) moving in two different directions (front-to-back and back-to-front) at three different speeds (times indicate how long it takes from the leading and trailing edges). Light gray horizontal bars at the bottom indicate stimulus presentation (n = 11 flies, 3 repetitions per fly).

We next determined the receptive field (RF) size of a single T3 neuron (**Figure 1E**) (Städele et al., 2020). We divided the right hemifield of the visual display into 10 azimuthal and 8 elevation rectangular sampling bins and presented flies with a 2.25° dark bar moving within each bin in orthogonal directions (see Methods). Responses to vertical and horizontal bar displacements were then multiplied to obtain the outer product (**Figure 1F**). To average RF size across flies, we selected the peak values of each bin and normalized them to the maximum value of the outer product. Finally, we spatially centered and averaged the RFs (**Figure 1G**). Average T3 RF size was mainly confined to the central 9° bin with almost no activity outside of it, consistent with a previous estimation (Tanaka and Clark, 2020). We tested directional selectivity by comparing responses to four cardinal directions. We found that T3 neurons were almost identically sensitive to leftward and rightward moving bars, as well as to upward and downward movements, although with a higher variability to the latter (**Figure 1H**).

Next, we explored how T3 respond to patterned “motion-defined” bars that robustly evoke saccades (Mongeau et al., 2019; Mongeau and Frye, 2017). A motion-defined bar is composed of the same random ON/OFF pattern as the stationary background, and therefore only detectable while in motion, by contrast to a classical luminance defined bar, which is brighter or darker than the surroundings and thus detectable even when stationary. Note that T4/T5 respond more strongly to a solid luminance-defined bar by comparison to a motion-defined bar (Keleş et al., 2018) due to sensitivity for longer spatial wavelength stimuli (optimum >= 15° wide solid bar) (Agrochao et al., 2020; Groschner et al., 2022; Gruntman et al., 2019; Shinomiya et al., 2019). Due to T3 rapid response kinetics to flicker (Keleş et al., 2020), we reasoned that these cells should be robustly excited by the motion-defined bars that drive saccades. We presented motion-defined bars moving front-to-back and back-to-front at three speeds. As expected, T3 neurons responded strongly to these stimuli by comparison to solid ON and OFF bars (**Figure 1I**). As bar speed increased the responses decreased monotonically (**Figure 1I**), both for motion-defined and for solid OFF and ON bars (**Supplementary Figure 1A**).

We coarsely assessed wide-field responses by T3 by presenting gratings of two different spatial frequencies moving at different velocities in two directions. As expected, wide-field responses showed phasic responses to individual cycles of the pattern, without selectivity for motion direction (**Supplementary Figure 1B**). Temporal frequency is the ratio of velocity to spatial wavelength, and thus if T3 behaves like T4/T5 then we would expect to observe responses tuned to the temporal frequency, not to the stimulus velocity. Oddly, the amplitude of responses to the λ=18° pattern moving at 2 Hz (36° s^-1^) were very similar to the responses to the λ=36° pattern moving at 1 Hz (36° s^-1^), indicating that response amplitude was tuned to stimulus velocity, not temporal frequency (Supplementary Figure 1c). Furthermore, the two spatial patterns presented at 90° s^-1^ (5 Hz and 2.5 Hz temporal frequency) produced identical amplitude responses (**Supplementary Figure 1C**).

As we did for T3, we also characterized the responses to bar stimuli by T2a neurons, which innervate layer 1, 2 and 9 of the medulla (**Supplementary Figure 2A**). Similar to T3, T2a presented a small RF and an omnidirectional sensitivity (**Supplementary Figure 2B-D**). T2a showed strong responses to luminance-defined bars (with a preference for ON transitions) moving at a low speed (18° s^-1^), but not at behaviorally relevant higher speeds (**Supplementary Figure 2E**). Moreover, they showed weak-to-no responses to motion-defined bars (**Supplementary Figure 2F**) and wide-field gratings (**Supplementary Figure 2G**). These results emphasize the distinct visual receptive field properties among different classes of columnar T-neurons, and distinguish potential behavioral importance of T3 neurons to the bar tracking behaviors of flies. We therefore focus our behavioral analysis on T3 neurons.

### Hyperpolarizing T3 reduces counter-directional object orientation by rigidly tethered flies

Rigidly tethered flies respond to a bar revolving around a cylindrical visual display (**Figure 2A**) with a compound counter-directional orientation response while the bar is in the visual periphery, switching to syn-directional tracking response as the bar approaches and crosses the visual midline (Reiser and Dickinson, 2010). **Figure 2A** shows, in schematic form, the direction that flies steer in response to a revolving bar. As the bar moves from the rear into the periphery, the fly initially steers toward the bar’s position, opposite its direction of motion (**Figure 2A**, lower). As the bar moves towards visual midline, the steering effort switches to track the direction of bar motion. The sum of the initial counter-directional and following syn-directional turns results in zero net steering effort once the bar is on visual midline - the fly is oriented directly at the bar. A defect in the counter-directional orientation phase would be expected to reduce the magnitude of the red trace, whereas a defect in syn-directional tracking would reduce the magnitude of the blue trace (**Figure 2A**, lower).

**Figure 2.**
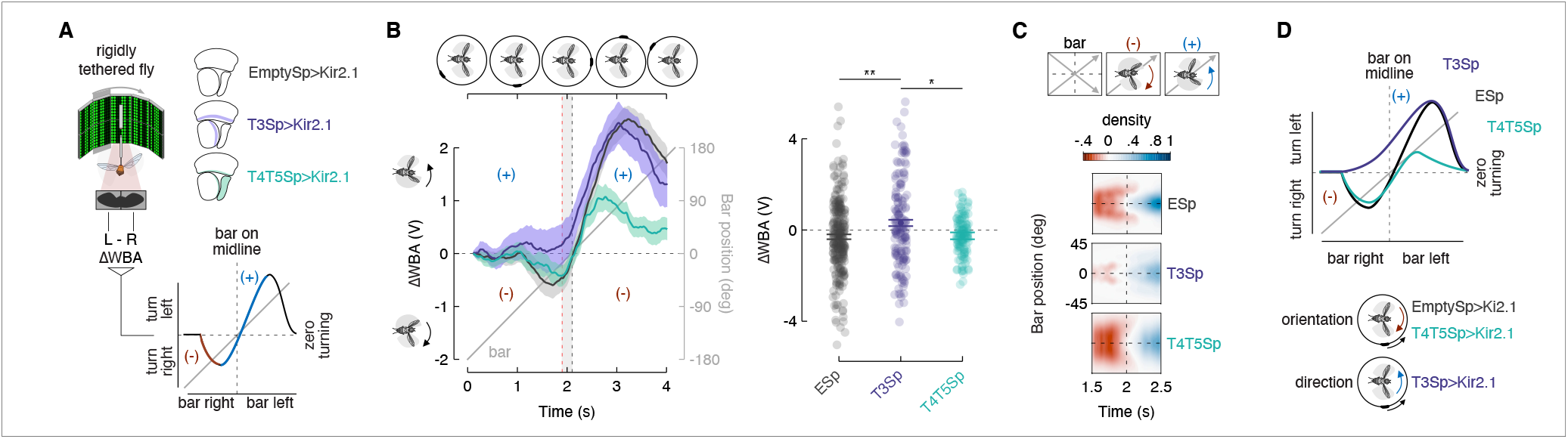
Constitutive silencing of T3 compromises the orientation response. (A) Left: cartoon of the rigid-tether setup in which a fly is glued to a tungsten pin and placed within a surrounding LED display presenting a random pattern of bright and bark stripes. An infrared diode above the fly casts a shadow on an optical sensor that records the difference between the left and right wing beat amplitudes (ΔWBA). Middle: schematic representation of the optic lobe regions where Kir2.1 channels were expressed in the three genotypes tested (data referred to T4/T5Sp>Kir2.1 flies are reproduced from Keleş et al., 2018). Bottom: schematic diagram of the experiment. A bar revolves around the fly (gray). Initially, steering is in the direction opposite bar motion plotted in red (-), followed by steering in the same direction as the bar plotted in blue (+). Depending on the strength of each response, the steering effort may be non-zero when the bar is at zero degrees (midline). (B) Top: schematic of the bar positions over time. Left: population average time series steering responses (mean ± s.e.m.) in the three genotypes tested (T4/T5Sp data replotted from Keleş et al., 2018) to a motion-defined bar revolving at 90° s-1 (responses to CCW rotations were reflected and pooled with CW responses). Gray shaded region (between the vertical red and black dashed lines) represents a 200 ms time window before the bar crosses the fly’s visual midline (n = 44 EmptySp>Kir2.1, n = 26 T3Sp>Kir2.1, n = 22 T4/T5Sp>Kir2.1). Note that T3Sp>Kir2.1 reduces counter-directional steering, whereas T4/T5Sp>Kir2.1 reduces syn-directional steering. Right: dot plot average ΔWBA values across the 200 ms time window per trial. Dark dots indicate the mean and the horizontal bars indicate s.e.m. A linear mixed model was used to fit the data and ANOVA with pairwise post-hoc comparisons adjusted using Bonferroni method were used to compare the three genotypes (F(2, 89) = 5.83, p = .004; EmptySp vs T3Sp: p = .005; EmptySp vs T4/T5Sp: p = 1; T3Sp vs T4/T5Sp: p = .03). (C) Heat maps of flies’ steering effort at the population level in the three genotypes as a function of the bar position (data are mirrored along the x-axis in order to get a uniform directional distribution). EmptySp and T4/T5Sp show a strong counter-directional response (red blob) while T3Sp show only a very slight counter-directional response. (D) Schematic summary of experimental results. T4/T5>Kir reduces the syn-directional tracking effort while leaving the counter-directional (-) orientation response intact. T3Sp>Kir2.1 reduces the counter-directional steering effort, leaving the syn-directional response (+) intact, and therefore steers leftward of the bar as it crosses midline.

Flies expressing outward cation channel Kir2.1 (Baines et al., 2001) in T4/T5 neurons showed weakened syn-directional tracking responses, but unaltered counter-directional orientation responses (Keleş et al., 2018) (**Figure 2B**), confirming that T4/T5 neurons are required for directional tracking, but not for positional orientation. We repeated this experiment after silencing T3 neurons. Compared to the responses of enhancerless split-Gal4 crossed with UAS-Kir2.1, the counter-directional orientation responses of T3 silenced flies were significantly reduced, whereas T4/T5 silenced flies were unaffected (**Figure 2B**). Flies with hyperpolarized T3 neurons responded to the revolving bar by steering (ΔWBA) always in the direction of motion. Thus, as the bar crossed visual midline, these animals showed a seemingly “anticipatory” response (ΔWBA > 0 at visual midline). By contrast, controls and T4/T5 silenced flies showed an initial counter-directional orientation response, followed by a weakened syn-directional phase superposing so that steering is balanced (ΔWBA = 0) as the bar crossed midline (**Figure 2B**). Quantification of the steering effort as the bar crossed midline indicates significant influence of hyperpolarizing T3, but not T4/T5 or the genetic control (**Figure 2B**, right).

To better visualize the finding that counter-directional orientation responses were strongly compromised by hyperpolarizing T3, we integrated the ΔWBA over time as a measure of optomotor wind-up. We then color coded each trace and zoomed in on trajectories passing near the visual midline. Reflecting and pooling responses to the two directions of bar revolution produced a heat map that reveals reduced orientation responses (red) for T3 silenced flies by comparison with controls and T4/T5 silenced flies (**Figure 2C**).

For bar stimuli that evoke significantly stronger syn-directional steering effort, such as a 30° wide solid dark bar, silencing T3 had no effect, whereas, as shown previously (Keleş et al., 2018), blocking T4/T5 neurons did (**Supplementary Figure 3A,B**). Confirming the results of **Figure 2**, the typical “anticipatory” response generated by winding up the optomotor system, which is dependent on T4/T5 neurons, was intact in T3 silenced flies (**Supplementary Figure 3C,D**). In summary, the syn-directional optomotor driven response to a moving bar is dependent upon T4/T5 activity, whereas the counter-directional orientational response is dependent upon T3 activity (**Figure 2D**).

### T3 hyperpolarization reduces saccadic bar tracking in magnetically tethered flies

In freely flying flies, orientation responses to visual objects trigger rapid body rotations called saccades for the functional analogy to our own gaze stabilizing rapid eye movements (Land, 1992; Tammero and Dickinson, 2002; van Breugel and Dickinson, 2012). Behavioral results and theoretical models suggest that both orientation responses by rigidly tethered flies and saccadic tracking responses by magnetically tethered flies could be coordinated by non directional (or omnidirectional) positional feature detectors with sensitivity to the high frequency transients generated by motion-defined bars, naturalistic stimuli that would not strongly activate directional motion detectors (Keleş et al., 2018; Mongeau and Frye, 2017; Reichardt and Poggio, 1976). T3 cells indeed show these physiological properties (**Figure 1**), and are required for intact counter-directional orientation steering effort (**Figure 2**). We therefore tested the functional role of T3 neurons for saccadic bar tracking by using magnetically tethered flies, free to steer in yaw on a frictionless pivot and execute robust body saccades (**Figure 3A**).

**Figure 3.**
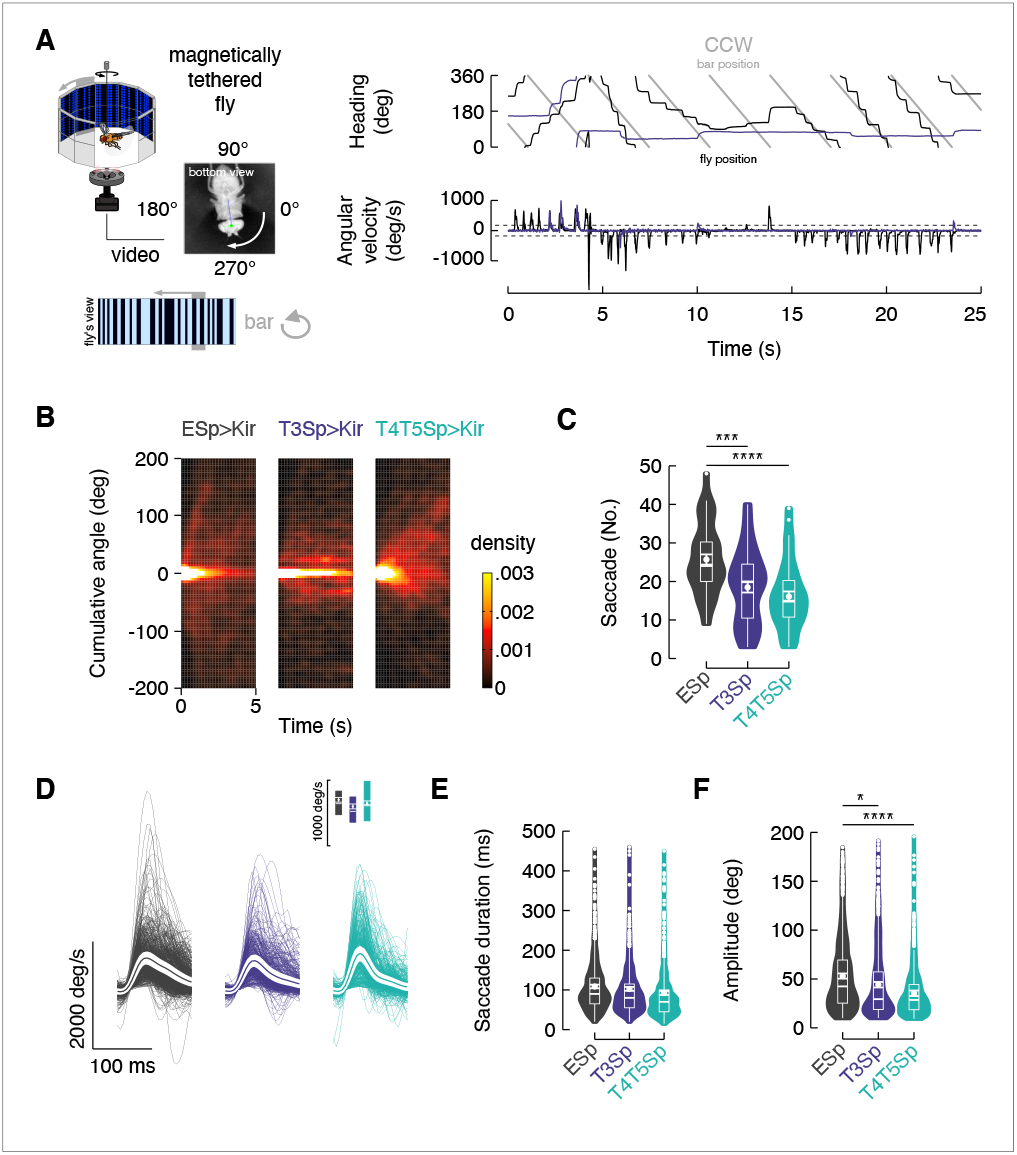
Flies with T3 hyperpolarized poorly track a motion-defined bar. (**A**) Top-left: cartoon of the magnetic-tether setup in which a fly is glued to a stainless steel pin and suspended within a magnetic field in turn placed within a surrounding LED display presenting a random pattern of bright and dark stripes. Infrared diodes illuminate the fly from below and a camera captures videos of the fly’s behavior from the bottom. Bottom-left: motion-defined bar moving CCW from the fly’s perspective. Top-right: wrapped heading traces from two representative flies (dark: EmptySp>Kir2.1; magenta: T3Sp>Kir2.1) responding to a CCW revolving motion-defined bar at 112.5° s^-1^ for 25 s. Gray line represents the bar position (in this plot it relates to the EmptySp fly). Bottom-right: filtered angular velocity profiles referring to the two representative flies at the top. Horizontal dashed lines indicate the threshold for detecting saccades. (**B**) Heat maps of the cumulative (i.e., unwrapped) angular distance traveled by flies within bins of 5 s in the three genotypes during the rotation of a motion-defined bar (n = 23 EmptySp>Kir2.1, n = 22 T3Sp>Kir2.1, n = 22 T4/T5>Kir2.1). (**C**) Violin-box plots of number of bar tracking saccades per trial in the three genotypes (pairwise post-hoc comparisons adjusted Bonferroni, EmptySp vs T3Sp: p = .0003; EmptySp vs T4/T5Sp: p < .0001; T3Sp vs T4/T5Sp: p = .55). Big white dots represent the mean, thin horizontal bars indicate s.e.m. and thick horizontal bars indicate the median. Small white dots on the violin tails represent outliers. (**D**) Average angular velocity (mean ± s.e.m.) during bar tracking saccades in the three genotypes. Top: peak angular velocity (pairwise post-hoc comparisons adjusted Bonferroni, EmptySp vs T3Sp: p = .40; EmptySp vs T4/T5Sp: p = 1; T3Sp vs T4/T5Sp: p = 1). Thin lines represent single saccades. (**E**) Violin-box plots of saccade duration (pairwise post-hoc comparisons adjusted Bonferroni, EmptySp vs T3Sp: p = .37; EmptySp vs T4/T5Sp: p = .13; T3Sp vs T4/T5Sp: p = 1). Central tendency measures as in **C**. (**F**) Violin-box plots of saccade amplitude (pairwise post-hoc comparisons adjusted Bonferroni, EmptySp vs T3Sp: p = .01; EmptySp vs T4/T5Sp: p = .001; T3Sp vs T4/T5Sp: p = 1).

Following the approach of prior work (Mongeau and Frye, 2017), we elicited bouts of tracking saccades by revolving a motion-defined bar against a stationary background (**Figure 3A**, right). In magnetically tethered flies, saccades are easily identified by characteristic impulses in angular velocity resulting in stepwise changes in flight heading (**Figure 3A**, right, **Video 2**). We hyperpolarized T3 and T4/T5 neurons by expressing Kir2.1 channels with split-Gal4 lines (**Video 3** and **4**). We first measured the cumulative angular distance that flies traveled in response to bar motion. Each trial was parsed into 5 second bins, normalized for initial heading (0°), and overlaid. Assuming bilateral symmetry, we reflected the CCW traces so as to have all traces representing responses to CW bar motion (**Figure 3B**, positive-going cumulative angle). We spatially pooled the overlaid traces to generate a heat map of the cumulative angular distance traveled by flies in each 5 second epoch. Empty vector controls dispersed within the first second, but T3 silenced flies remained concentrated at their initial heading, indicating that they were not tracking the motion-defined bar (**Figure 3B**). T4/T5 silenced flies dispersed similar to controls. We next enumerated body saccades per trial (Mongeau and Frye, 2017). Both T3 and T4/T5 silenced flies performed fewer tracking saccades per trial than controls, suggesting dependence of both of these cell types for triggering saccades (**Figure 3C**). We measured the angular velocity profile of saccades, which did not differ across the three genotypes (**Figure 3D**), and same for saccade duration (**Figure 3E**). However, saccade amplitude was slightly reduced in T3 blocked flies and strongly reduced in T4/T5 blocked flies (**Figure 3F**). As expected, in response to rotation of the full wide-field panorama, T4/T5 silenced flies showed reduced smooth tracking gain (ratio of body rotation to stimulus rotation, **Supplementary Figure 4A,B**) due to compromised directional optomotor responses (Bahl et al., 2013).

### Inducible T3 depolarization enhances saccadic bar tracking in magnetically tethered flies

Constitutive hyperpolarization of T3 shows that normal excitatory activity in these cells is required for saccadic bar tracking pursuit in flight (**Figure 3**). We sought to further support this result with an inducible perturbation. We expressed CsChrimson (Klapoetke et al., 2014) channels in T3 and T4/T5 neurons and stimulated with periods of CsChrimson-activating red light intermittently OFF and ON with 5 second intervals, while presenting flies with a revolving motion-defined bar (**Figure 4A**). This approach activates the population of columnar neurons together, rather than in the spatially localized retinotopic manner they would be normally active. Therefore, we consider this approach to function as a loss-of-function perturbation, rather than a gain-of-function excitation of T3, and is of course dynamic rather than static (e.g., Kir2.1). Our aim was to assess how bar tracking behavior was impacted by either phase of the perturbation: saturating ensemble depolarization, or recovery. In controls, the light ON did provoke small changes in the flies’ heading (**Figure 4B**, top black). Activation of T3 neurons at high LED intensity tended to reduce or eliminate active bar tracking, instead provoking seemingly random exploratory saccades (**Figure 4B**, middle magenta). Ensemble depolarization of T4/T5 evoked seemingly stronger bar tracking (**Figure 4B**, bottom green). To visualize the population effects of dynamic depolarization, we binned together the unwrapped heading traces segregated by OFF and ON epochs and normalized to the initial heading, and generated heat maps of the resultant cumulative heading angle. We split the heat maps into two spatial windows, initial responses 0-200° of cumulative angle, and later responses of 200-500° range. For the initial accumulation of heading change, up to 200°, the heat maps between CsChrimson OFF and ON epochs were not obviously different across the three genotypes (**Figure 4C**, lower). However, light-gated depolarization of T3 neurons caused cessation of bar responses prior to a single 360° revolution around the circular arena, whereas controls and T4/T5>CsChrimson flies, on average, continued to steer throughout the trail (**Figure 4C**).

**Figure 4.**
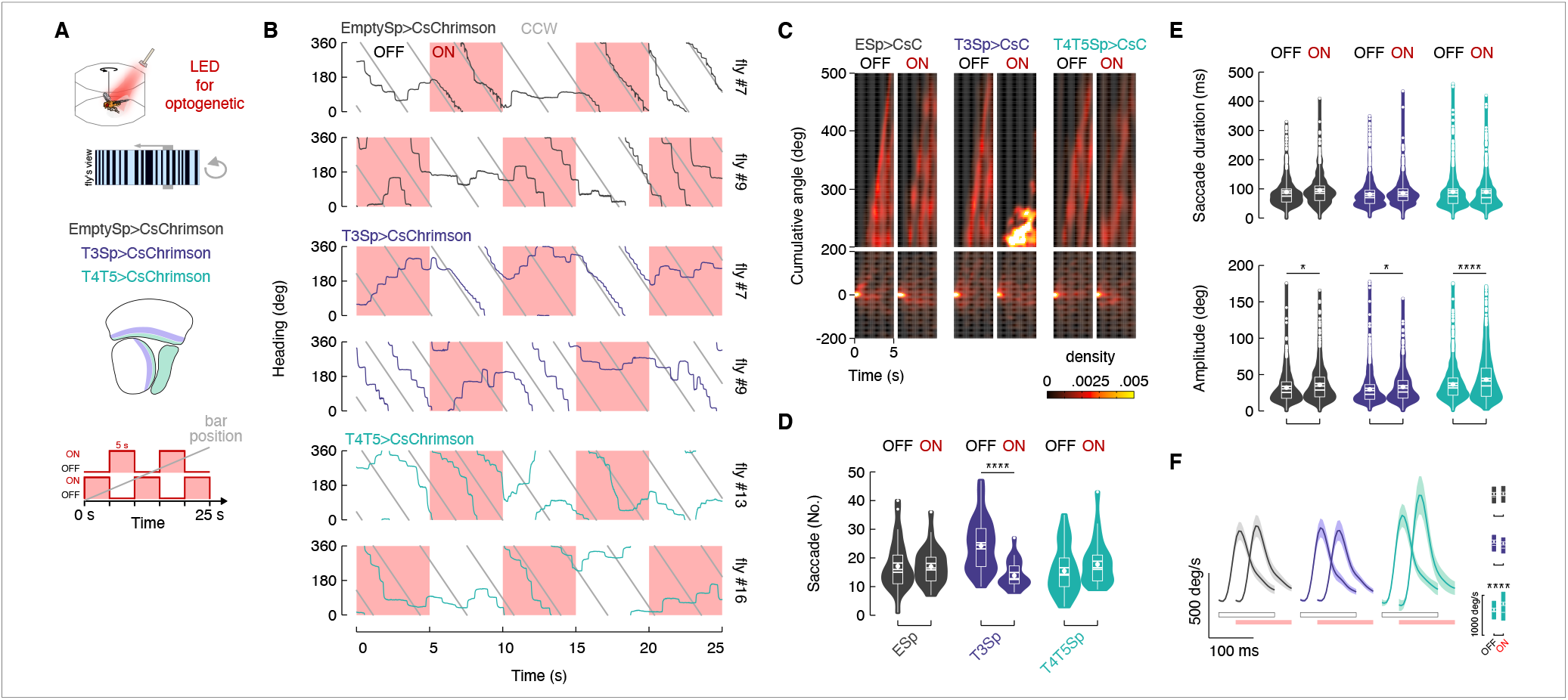
Depolarization of T3 increases bar tracking saccades. (**A)** Top: cartoon of the magno-rigid setup implemented with an optogenetic LED for cells’ stimulation. Middle: genotypes tested and schematic representation of the optic lobe regions innervated by T3 and T4/T5. Bottom: schematic representation of the stimulation protocol: repetition of 5 s optogenetic LED on followed by 5 s LED off for 25 s. On/off starting was randomly selected. (**B**) Wrapped heading traces from six representative flies (dark: EmptySp>CsChrimson; magenta: T3Sp>CsChrimson; green: T4/T5Sp>CsChrimson) responding to a CCW revolving motion-defined bar as in **Figure 3A**. Red shaded regions represent periods of LED on. EmptySp>CsChrimson and T4/T5EmptySp>CsChrimson flies were slightly affected by the LED on while T3Sp>CsChrimson flies stop chasing the bar or start to turn CW. (**C**) Heat maps of the cumulative angular distance traveled by flies within bins of 5 s in the three genotypes during the rotation of a motion-defined bar (n = 20 EmptySp>Kir2.1, n = 22 T3Sp>Kir2.1, n = 21 T4/T5>Kir2.1). The map was divided in two windows to highlight the late component of the response where T3Sp>CsChrimson flies remain stuck during the stimulation periods. (**D**) Violin-box plots of number of bar tracking saccades during the periods of on and off stimulations (pairwise post-hoc comparisons adjusted Bonferroni, EmptySp: p = 1; T3Sp: p < .0001; T4/T5Sp: p = 1). (**E**) Top: violin-box plots of saccade duration between on and off stimulations (pairwise post-hoc comparisons adjusted Bonferroni, EmptySp: p = .19; T3Sp: p = .84; T4/T5Sp: p = 1). Bottom: violin-box plots of saccade amplitude by stimulation condition (pairwise post-hoc comparisons adjusted Bonferroni, EmptySp: p = .02; T3Sp: p = .03; T4/T5Sp: p < .0001). (**F**) Left: average angular velocity (mean ± s.e.m.) of saccades. Light gray and red horizontal bars at the bottom indicate the stimulation conditions (red: LED on; gray: LED off). Right: box plots of peak angular velocity in the on and off LED conditions (pairwise post-hoc comparisons adjusted Bonferroni, EmptySp: p = 1; T3Sp: p = .05; T4/T5Sp: p < .0001).

We next assessed how saccades were triggered under CsChrimson perturbation. T3>CsChrimson flies showed on average a dramatic increase in the number of saccades during the OFF epochs compared to the ON epochs, whereas the number of saccades was similar for T4/T5 activated flies and Empty>CsChrimson controls (**Figure 4D**). These results indicate that, in agreement with the cumulative steering angle results, population depolarization of T3 neurons perturbed this visual pathway, reducing bar-evoked saccades. Yet, when the optogenetic stimulus was switched OFF, recovery from sustained depolarization strongly revived saccadic bar tracking in flies expressing CsChrimson in T3 (**Figure 4D**).

We next tested whether saccade dynamics were affected by optogenetic stimulation of T3. Saccade duration was affected neither by CsChrimson expression nor optical condition in any genotype (**Figure 4F**). Saccade amplitude increased modestly in all genotypes likely due to an artifact of the red light (Klapoetke et al., 2014; Wu et al., 2016), but in T4/T5 this effect was pronounced (**Figure 4E**). Without any change in saccade duration, the increase in saccade amplitude in T4/T5 activated flies yielded a correspondingly strong increase in saccade angular velocity only for T4/T5>CsChrimson (**Figure 4F**). In controls and T3 activated flies the saccade angular velocity was similar for OFF and ON optogenetic stimulation epochs. These results indicate that although normal T3 function is required for triggering saccades to track moving objects (**Figure 3** and **4B-D**), T3 neurons are not involved in controlling saccade amplitude (**Figure 4E**). By contrast, normal T4/T5 function is essential to controlling saccade amplitude but not in triggering object tracking saccades (**Figure 3F** and **4E,F**).

Flies execute dynamically distinct classes of saccades for tracking objects, for avoiding objects, and for minimizing wide-field perturbations (Mongeau et al., 2019; Mongeau and Frye, 2017). To test whether T3 signals are used specifically for object tracking saccades, we tested flies with a rotating wide-field panorama, no bar. Our prediction was that if T3 functions specifically for bar tracking saccades, then silencing them should have no influence over wide-field evoked saccades, and *vice versa* for T4/T5 neurons. In support of this prediction, the number of wide-field optomotor saccades in T3>CsChrimson flies did not change significantly between OFF and ON epochs but increased robustly for T4/T5>CsChrimson flies (**Figure 5A**). Saccade duration and amplitude were unaffected for any genotype or optical activation condition (**Figure 5B,C**). However, the angular velocity of wide-field evoked saccades increased for T4/T5 activated flies during ON epochs (**Figure 5D**). Taken together, our results show that T4/T5>CsChrimson facilitates more optomotor saccades with higher velocity in response to both a small-field object and wide-field panorama, whereas the effects of T3>CsChrimson are specific to object tracking saccades.

**Figure 5.**
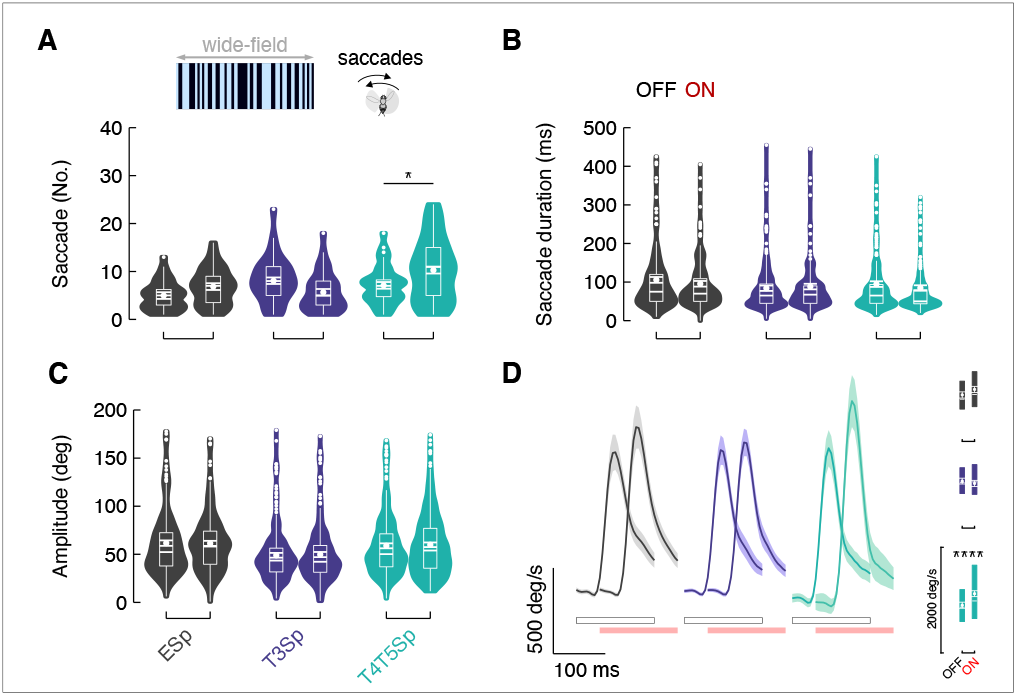
Depolarization of T4/T5 increases optomotor saccades. (**A**) Top: representation of the wide-field pattern of bright and dark random stripes rotating around the fly. Bottom: violin-box plots of number of optomotor saccades by stimulation condition (pairwise post-hoc comparisons adjusted Bonferroni, EmptySp: p = .89; T3Sp: p = .19; T4/T5Sp: p = .02). (**B**) Optomotor saccade duration between on and off stimulations (pairwise post-hoc comparisons adjusted Bonferroni, EmptySp: p = .32; T3Sp: p = 1; T4/T5Sp: p = .08). (**C**) Optomotor saccade amplitude by stimulation condition (pairwise post-hoc comparisons adjusted Bonferroni, EmptySp: p = 1; T3Sp: p = 1; T4/T5Sp: p = 1). (**D**) Left: average angular velocity (mean ± s.e.m.) of optomotor saccades. Light gray and red horizontal bars at the bottom indicate the stimulation conditions (red: LED on; gray: LED off). Right: box plots of peak angular velocity in the on and off LED conditions (pairwise post-hoc comparisons adjusted Bonferroni, EmptySp: p = 1; T3Sp: p = 1; T4/T5Sp: p < .0002).

### T3 calcium dynamics support an integrate-and-fire model for triggering saccades

Prior work has shown that bar-tracking saccades are triggered neither by absolute retinal position, nor by bar velocity, but rather by a threshold in the spatial integral of bar position over time (Mongeau and Frye, 2017). How could T3 contribute to this computation? For bar tracking saccades, the product of time (inter-saccadic interval, ISI) and angular distance traveled by the bar (pre-saccade error angle) is invariant across bar speed (Mongeau and Frye, 2017), and thus the terms are inversely proportional. When the bar moves fast, it generates a large pre-saccade error angle, a saccade is triggered early, and the inter-saccadic interval is short. When the bar moves slowly, the error angle is small, and the inter-saccadic interval is extended (**Figure 6A**). These spatial dynamics place constraints on T3 action, since the number of retinal facets and corresponding neural columns and T3 neurons that are stimulated by object motion are proportional to object speed, and inversely related to the visual dwell time on each facet, parameters that highlight T3 function for triggering saccades.

**Figure 6.**
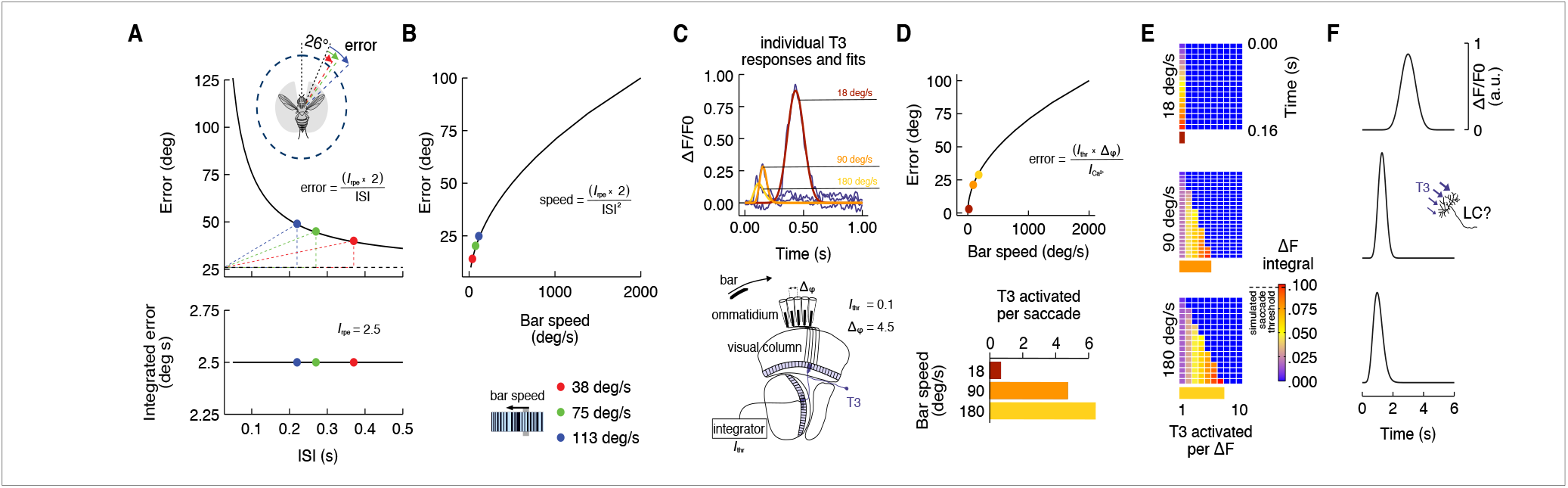
Integrate-and-fire model physiologically inspired on T3 calcium dynamic. (**A**) Top: pre-saccade error angles modeled by using the inverse function of the integrated error (Mongeau and Frye, 2017). Knowing the inter-saccadic intervals (ISI) from previous behavioral experiments as a function of the bar speed and the integrated retinal position error (*I*_*rpr*_), we can compute the pre-saccade error angle as a function of ISI. Modeled error angles are very close to the average errors from behavioral experiments (ISI = 0.22 s, Error = 49°; ISI = 0.27 s, Error = 45°; ISI = 0.37 s, Error = 40°). Dashed black line represents the threshold from which the integration of the bar position over time starts (26°). The area of the triangles defined by the dashed colors lines is constant and represents the *I*_*rpr*_. Bottom: integrated retinal position error (*I*_*rpr*_ = 2.5° s) derived from previous experiments work (Mongeau and Frye, 2017) that represents the multiplication of pre-saccade error angle by ISI. (**B**) Modeled pre-saccade error angle as a function of the bar speed. Higher speeds require larger pre-saccade error angles. (**C**) Top: average calcium responses of T3 neurons (magenta) to a moving motion-defined bar at three different speeds (data from **Figure 1I**). These responses were fitted using nonlinear regression analysis (yellow: 180° s^-1^; orange: 90° s^-1^; red: 18° s^-1^). Areas under the curves were computed (*I*_*Ca2+*_). Bottom: schematic representation of the fly visual lobe with a 1D organization of T3 neurons that are sequentially activated by a moving bar across the retina. A T3 neuron is present in each column and the interommatidial angle (Δ_φ_) is ∼ 4.5°. We set an arbitrary integrated threshold (*I*_*thr*_ = 0.1) that a downstream partner of T3 would use to trigger a saccade. (**D**) Top: modeled pre-saccade error angles based on *I*_*Ca2+*_ for different speeds (Δ_φ_ is known and *I*_*thr*_ is arbitrary). These error values scale exactly like the modeled behavioral pre-saccade errors as a function of speed. Bottom: number of T3 cells that in a 1D space would be required to trigger a saccade according to the model. (**E**) Space-time plots of *I*_*thr*_ accumulating over time across cells depending on the bar speed. (**F**) Simulated responses by speed of a hypothetical integrator downstream of T3 neurons. LC17 cells might play this role.

Prior behavioral experiments (Mongeau and Frye, 2017) indicated that the fly’s desired ‘set point’ or reference position of the bar is not visual midline, but rather is offset laterally approximately 26° (**Figure 6A**, inset). If the bar moves quickly away from the reference position, then many retinal facets would be stimulated with little dwell time, whereas if the bar moves slowly, fewer facets would be stimulated but with larger dwell time on each. For a downstream integrating neuron, short dwell time corresponds to a low amplitude signal for a given columnar input, requiring many such signals to charge the integrator to threshold. If the bar moves slowly, larger amplitude signals mean fewer are needed to reach firing threshold. This scheme requires neural responses that are inversely proportional to bar speed, which we observed in T3 (**Figure 1I**): low amplitude calcium responses for fast moving bars, and large responses for slow moving bars.

Do T3’s calcium responses scale to the behavioral data (**Figure 6B**) and support a spatiotemporal integrate-and-fire model? To test this hypothesis, we developed a simple physiological model for spatiotemporal integration. We fitted the curves of the GCaMP responses by using a nonlinear least squares method which allowed us to estimate the model parameters and to calculate the respective integrals (*I*_*Ca2+*_) (**Figure 6C**, top). We used these fits in a model 1D cell array corresponding to the fly’s horizontal visual midline (**Figure 6C**, bottom). An arbitrary threshold (*I*_*thr*_ = 0.1) represents the integrated retinal position error and, since there is one T3 cell per medulla column (Takemura et al., 2015), we set the interommatidial angle (Δϕ) = 4.5°. By using a simple equation where we divided the *I*_*th*_ by *I*_*Ca2+*_ and then multiplied by Δϕ (**Figure 6D**, inset), we computed the retinal position error at which flies would trigger a saccade based on the modeled T3’s calcium responses (**Figure 6D**). The physiologically-inspired retinal position error fits very well on the curve representing the behavioral relationship between retinal error and bar speed (**Figure 6D**). This allowed us to calculate the number of columnar neurons that would have been stimulated prior to triggering a saccade for each stimulus velocity (**Figure 6D**, lower). Depending on the bar speed (hence the dwell time within each columnar cell’s RF), the columnar activation of the cell array would charge a downstream integrator to firing threshold for triggering a saccade (**Figure 6E,F**). A simulated fly’s position based on this physiologically-inspired control system recapitulates quite well the real fly behavior (**Supplementary Figure 5**). Our simple model provides a parsimonious explanation for how T3 neurons can provide behaviorally relevant signals to trigger object tracking saccades.

## Discussion

Like virtually every animal with image forming eyes, including humans, flies use smooth optomotor movements to stabilize gaze and to maintain visual course control during locomotion (Land, 1992). It is widely accepted that optomotor stabilization reflexes in flies are elicited by patterns of wide-field optic flow, which is detected by spatially integrating the signals from two identified classes of directionally selective motion detecting neurons, T4 and T5, columnar neurons with narrow receptive fields (small-field) that sample the entire visual field (Mauss and Borst, 2020; Yang and Clandinin, 2018). T4/T5 neurons innervate the lobula plate, where they synapse with wide-field collating neurons such as the horizontal system class (Shinomiya et al., 2022) of lobula plate tangential cells (LTPCs). LPTCs have complex directional receptive fields that act as spatial filters for the patterns of optic flow generated by specific flight maneuvers, and which in turn coordinate smooth optomotor responses (Busch et al., 2018).

In parallel with smooth continuous optomotor movements, flies, like humans, also execute saccades to shift gaze, both during walking (Geurten et al., 2014) and in flight to track or avoid salient moving objects (Mongeau et al., 2019; Mongeau and Frye, 2017). Models of smooth optomotor visual stabilization do not account for saccadic object pursuit (Fenk et al., 2014; Reichardt and Poggio, 1976), which instead are believed to be coordinated by a pathway operating in parallel to T4/T5 directional motion detectors (Aptekar et al., 2012; Bahl et al., 2013). Here we provide evidence for the identity of this parallel pathway; columnar T3 neurons arise from the same optic ganglion as T4/T5, but rather than innervating the lobula plate center for motion vision, T3 terminate in the outer lobula, a center of visual feature detection (Keleş and Frye, 2017a). To date, what is known of T3 cholinergic chemical synapses (Konstantinides et al., 2018) includes lobula columnar projection neurons LC11 and LC17 (Tanaka and Clark, 2022). LC11 is a small object movement detector that plays no role in flight control, but rather seems to coordinate conspecific social interactions (Ferreira and Moita, 2020; Keleş and Frye, 2017b). A behavioral screen of freely walking flies showed that optogenetic activation of LC17 induced turning responses (Wu et al., 2016), and LC17 has been shown to respond to looming stimuli (Klapoetke et al., 2022), but any potential role in saccadic object tracking is unknown.

The receptive field properties of T4/T5 are well suited to control smooth, directional optomotor responses, whereas T3 are better suited to detect the features of visual objects that flies track. For example, T4 and T5 are half-wave rectified for contrast increments and decrements, respectively, and are tuned to the temporal frequency of a moving pattern (Joesch et al., 2010). By contrast, T3 are distinguished from T4/T5 (and also from neighboring T2a neurons) by full wave rectification (Keleş et al., 2020) (i.e., ON-OFF selectivity), omnidirectionality, and broad sensitivity to temporal frequency sensitivity (Figure **1** and Supplementary Figure 2). T3 responds vigorously to the high frequency (finely textured) stimuli that robustly drive object tracking behavior in flight (**Figure 1**), whereas these same object stimuli drive T4/T5 to a significantly lesser extent (Keleş et al., 2018).

To our knowledge, this is the first identification of a visual neuron type that specifically serves saccadic object tracking behavior (Mongeau et al., 2019; Mongeau and Frye, 2017). However, T3 does not act alone in this capacity. Our evidence suggests that T4/T5 coordinates distinct components of saccadic tracking behavior, possibly via identified projection neurons that interconnect the lobula plate and deeper layers of the lobula (Shinomiya et al., 2022). A control theoretic model that accounts well for the properties of saccadic object tracking requires two key computations: (1) an integrate- and-fire threshold operation to trigger a saccade, which depends upon the accumulation of retinal error, and (2) a torque magnitude variable that regulates saccade amplitude, which depends upon object direction and speed (Mongeau and Frye, 2017). Our evidence suggest that T3 neurons participate in triggering saccades, because their synaptic perturbation suppresses both orientation steering effort by the wings (**Figure 2**) and object pursuit saccades (**Figure 3** and **4**), while a model based on the integrated output of T3 neurons captures the spatiotemporal threshold dependence of bar tracking behavior (**Figure 6**). By contrast, our evidence shows that directional motion detectors T4 and T5 coordinate saccade amplitude, because constitutive synaptic blockade reduces saccade amplitude (**Figure 3**), and depolarization increases saccade amplitude (**Figure 4**). Finally, T3 perturbation does not influence the control of wide-field evoked saccades (but silencing T4/T5 does), an important result that highlights T3 selective role in object tracking saccades (**Figure 5**). Our working model is no doubt an incomplete accounting of all of the underlying control circuit interactions. Nevertheless, we have for the first time provided a strong conceptual framework, and identified its key elements, for the parallel neural control of smooth optomotor stabilization and saccadic object tracking control systems within a key model system.

## Methods

### Fly Strains

For all experiments we used 3-5 days old female Drosophila melanogaster reared on standard cornmeal molasses at 25°C, 30%-50% humidity entrained to 12 h light/12 h dark cycle. All behavioral experiments involving silencing of targeted neurons were performed with flies carrying at least one wild-type *white* allele. We were not blind to genotype. Fly lines and their origins are listed in Supplementary Table 1 and 2.

### Imaging setup

Two-photon calcium imaging was performed on a modified upright microscope (Axio Examiner, Zeiss), exciting the specimens with a Ti:Sapphire laser (Chameleon Vision, Coherent) tuned to 920 nm (power at the back aperture ∼ 25 mW). We imaged with a 20x water-immersion objective (W Plan-Apochromat 20x/1.0 DIC, Zeiss). Data acquisition was controlled by Slidebook (version 6, 3i). Single plane images were taken at ∼ 10 Hz with a spatial resolution of approximately 285 × 142 pixels (100 × 50 μm, pixel size ≅ 0.35 μm, dwell time ≅ 2.5 μs). GCaMP6f responses were recorded from the presynaptic terminals of T3 neurons. For each preparation, we identified the most caudal presynaptic terminals and then shifted the ROI downwards ∼ 30 μm. Images and external stimulations were synchronized *a posteriori* using frame capture markers (TTL pulses output from Slidebook) and stimulus events (analog outputs from the LED display controller) sampled with a data acquisition device (DAQ) (PXI-6259, NI) at 10 kHz. The DAQ interfaced with MATLAB (R2020a, MathWorks) via rack-mount terminal block (BNC-2090, NI).

### Fly preparation for imaging

Flies were cold anesthetized at ∼ 4°C by using a thermoelectric cooling system and mounted on a custom 3D printed fly holder (Weir and Dickinson, 2015). Specifically, flies were gently pushed through a hole etched on a stainless-steel shim so that the dorsal thorax protruded from the dorsal aspect of the horizontally mounted shim and the ventral thorax with the abdomen remained below. The head was pitched forward to expose its posterior surface without stretching the neck connective. Flies were secured using UV-curable glue (44600, Dreve Fotoplast Gel) around the posterior-dorsal cuticle of the head capsule, and across the dorsal thorax. To reduce movement artifacts during the recordings, we immobilized the legs and the proboscis with low melt point bees-wax (Waxlectric-1, Renfert). We used fine forceps (Dumont #5SF, Fine Science Tools) to remove the posterior cuticle, fat bodies and post-ocular air-sac obstructing the view of the right optic lobe. We severed muscles 1 and 16 to reduce brain movements (Demerec, 2008). The brain was bathed in physiological saline containing (in mM): 103 NaCl, 3 KCl, 1.5 CaCl_2_, 4 MgCl_2_, 26 NaHCO_3_, 1 NaH_2_PO_4_, 5 N-Tris (hydroxymethyl) methyl-2-aminoethane-sulfonic acid (TES), 10 trehalose, 10 glucose, 2 sucrose. The saline was adjusted to an osmolarity of 273-275 mOsm and a pH of 7.3-7.4. The brain was continuously perfused with extracellular saline at 1.5 ml/min via a gravity drip system and the bath was maintained at 22°C by an inline solution heater/cooler (SC-20, Warner Instruments) and temperature controller (TC-324, Warner Instruments).

### Visual stimuli for imaging

During two-photon imaging, visual stimuli were presented on a LED display (Reiser and Dickinson, 2008) composed of 48 panels arranged in a semi-cylinder (Panels arena, IO Rodeo). The display covered ±108° in azimuth and ±32° in elevation. Each LED subtended a visual angle of ∼ 2.25°. To reduce the light intensity from the LED display, three layers of blue filter (R59-indigo, Rosco) were placed over the display. The display had, at its maximum intensity, an irradiance of ∼ 0.11 μW m^-2^ (recorded at the fly’s position) at the spectral peak of 460 nm (full width at half maximum: 243 nm). Visual patterns were generated and controlled using custom-written MATLAB scripts that communicated to a custom designed controller (IO Rodeo) via a serial port, which in turn communicated to the panels via a rapid serial interface (https://reiserlab.github.io/Modular-LED-Display/Generation%203/). To account for the angle the fly’s head had when mounted on the holder, the display was tilted 30° from the horizontal plane. We recorded from the right optic lobe, and the stimulus coordinates are referred to the fly’s head position (Figure 1c). Therefore, azimuthal and elevation position of the stimuli are centered to the fly’s visual equator and prime meridian.

### Speed tuning

Visual stimulation was confined to the right half of the visual field, ipsilateral to the recording site. Due to the far peripheral blind spot generated by the imaging stage, visual stimuli were presented within a restricted window of 72° x 72° with the left edge abutting the visual prime meridian. The brightness of the display background, outside the stimulus window, was set to 50% maximum. The stimulus set was presented in random block design, repeated 3 times. Each visual stimulation lasted 7.5 s and was composed by 0.5 s of uniform background (50% maximum intensity), 0.5 s of static pattern onset within the stimulation window, variable duration (depending on the stimulus speed) visual motion (maximum 4.5 s) followed by 2 s lingering static pattern. After each visual stimulation, 2 s of rest with the display off (0% of maximum intensity) interspersed the trials to prevent adaptation.

### Receptive field mapping and directional selectivity

As previously described (Städele et al., 2020), to characterize the functional receptive field (RF) of T3 and T2a neurons, we computed the integrated responses to a 2.25° x 72° (width x height) dark bar (single pixel stripe) moving horizontally with the responses to a 72° x 2.25° dark bar moving vertically. The single pixel dark stripe moved within a 9° spaced bin (4 pixels) in all four cardinal directions (upward, downward, leftward and rightward) at 18° s-1, allowing us to extract directional selectivity information. Sweep trials of different directions were randomly presented over a region of the visual field between 0° and 90°in azimuth and between −36° and +36° in elevation. We divided this region in 10 horizontal bins (each one of 9° x 72°) and 8 vertical bins (each one of 90° x 9°). Each sweep (36 in total) was composed of 6 s of rest period with a uniform bright background (50% of maximum intensity) and 0.5 s of stripe motion within a bin. Since we did not observe differences in the directional selectivity of T3 and T2a for horizontal or vertical sweeps, leftward and rightward movement responses were averaged, as well as downward and upward responses. To assess the RF size, these averaged time series for the 10 horizontal bins and the 8 vertical bins were multiplied together, obtaining a matrix of activity (Figure 1f). The matrix for each fly was then simplified by taking the values of activity peak and spatially normalized to the bin showing the maximum activity peak (RF center). Finally, the peak values were normalized to the value at the center of the RF (bin #0). We averaged the normalized matrices for each fly to obtain a heatmap of the RF for T3 neurons (**Figure 1G**). The directional selectivity was estimated by considering the activity peaks of the RF center for each sweep separately and plotting them in a polar plot (**Figure 1H**). Flies whose anteroposterior imaging plane moved during the experiment were no longer considered for the analysis.

### Imaging data analysis

Image stacks were exported from Slidebook (™) in (16-bit) .tiff format and imported into MATLAB for analysis. A user-friendly custom toolbox developed by Ben J. Hardacastle (https://github.com/bjhardcastle/SlidebookObj) allowed us to correct the images for motion artifacts along the x-y plane using a DFT-based registration algorithm (Guizar-Sicairos et al., 2008) and to manually draw a region of interest (ROI) around the presynaptic terminal of an active neuron located approximately in the middle of the lobula. We opted for a manual identification of the ROI to specifically consider morphology and position of the neuron investigated. Through this approach, combined with the consistent z-position of our imaging plane, we were able to identify T3 neurons across preparations at the same location in the neuropile, exhibiting similar spatial receptive fields. A time-series was generated by calculating the mean fluorescence intensity of pixels within the ROI in each frame (F_t_). These mean values were then normalized to a baseline value as ΔF/F = (F_t_ – F_0_)/F_0_, where F_0_ was the mean of F_t_ during the 0.5 s preceding stimulus onset. To compute average time-series across preparations with small variations in TTL synchronization, traces were resampled using linear interpolation. This procedure did not cause any detectable change in the original data.

### Rigid-tether setup

In the rigid tether paradigm, flies were cold anesthetized at ∼ 4°C and tethered to tungsten pins (diameter: 0.2 mm) using UV-curable glue (Watch Crystal Glue Clear). The pin was placed on the dorsal thorax so as to get a final pitch angle of the fly (between 35° and 45°) similar to body angle and wing stroke plane during free hovering (Fry et al., 2003). Before running the experiment, flies were left to recover upside-down in a custom designed pin holder, in turn placed into a covered acrylic container, for ∼ 30 min at room temperature (∼ 22°C). Inside the container, we put a small bowl filled with water to maintain humidity and avoid desiccation. To reduce flight energy expenditure, as soon as flies recovered from the anesthesia and started flying, we gently offered them a small square piece of paper (Kimwipes, Kimberley-Clark) ∼ 3 mm side, that they generally clung to with their legs without flying. Flies were then positioned in the center of a cylindrical LED panel display (Reiser and Dickinson, 2008) that covered ±165° in azimuth and ±47° in elevation. The display was composed of 96(h) x 32(v) LEDs (emission peak: 568 nm) each LED subtending 3.75° on the fly’s retina. Flies were illuminated from the top with an infrared diode (emission peak: 880 nm) which cast a shadow of the beating wings onto an optical sensor. An associated “wing-beat analyzer” (JFI Electronics Laboratory, University of Chicago) converted the optical signal into an instantaneous voltage measuring right and left wing beat amplitude (WBA) and frequency (WBF). The difference in the left and right WBA (ΔWBA), which is highly correlated with the fly’s steering effort in the yaw axis (Tammero et al., 2004), connected to the panel display controller to close a feedback loop with the rotational velocity of the visual display. Signals from the wing-beat analyzer and from the panel display controller, encoding the visual display position, were recorded on a DAQ (Digidata 1440A, Molecular Devices) at 1 kHz. The data acquisition was triggered through a voltage step sent by a second DAQ (USB-1208LS, Measurement Computing) interfaced with MATLAB that in turn controlled the pattern presentations. For silencing experiments, flies expressing Kir2.1 tagged with green fluorescence protein (GFP) were dissected after the behavioral recordings under a fluorescence stereomicroscope (SteREO Discovery.V12, Zeiss) to confirm the expression in the targeted cells.

### Rigid-tether visual stimuli

Each trial was composed of 5 s in closed-loop with a bar and a variable time in open-loop test depending on the speed of the stimuli presented to the fly. The closed-loop periods ensured the fly was engaged in the task, while the open-loop periods were tested for responses to the visual stimuli. A full set of stimuli was randomized and repeated 3 times, with a total duration of ∼ 5 min. Flies that stopped flying during the experiment or that failed to frontally fixate a dark bar on a uniform bright background during a pre-experiment assessment period, were not included in the analysis. All stimuli used in these experiments had patterns of bright (100% of maximum intensity) and dark pixels (0% of maximum intensity). The luminance-defined bar represented a 30° x 94° (width x height) dark bar moving on a uniform bright background. The motion-defined bar represents a 30° x 94° bar of random bright and dark vertical stripes moving over an analog static background of randomly distributed bright and dark stripes. This pattern was generated using a custom-written MATLAB code as previously described (Keleş et al., 2018). Briefly, the random-stripes patterns had an equal number of bright and dark pixels and a high-pass filter ensured that no bright or dark contiguous stripes exceeded 22.5° in width (6 stripes). A random-stripes pattern was randomly picked up, every time it was needed, from 96 different options to avoid pattern-specific behavioral artifacts. Clockwise (CW) and counter-clockwise (CCW) directions were always considered in bar revolving experiments. However, assuming bilateral symmetry, we reflected the time series responses to counter-clockwise (CCW) stimuli and pooled them with clockwise (CW) responses.

### Magnetic-tether setup

In the magnetic tether paradigm, flies were cold anesthetized as for the rigid tether paradigm and glued to stainless steel pins (diameter: 0.1 mm, Fine Science Tools) as previously described (Bender and Dickinson, 2006; Duistermars and Frye, 2008; Mongeau and Frye, 2017). Briefly, pins were trimmed to be 1 cm length and placed on the dorsal thorax in order to get a final fly’s pitch angle of approximately 30°. Flies were then left to recover for ∼ 30 min upside-down sticking out of a polystyrene block. To reduce flies’ fatigue, as done for the rigid tether paradigm, flies were provided with a small piece of paper. The visual stimulation was performed using a cylindrical LED panel display covering 360° in azimuth and 56° in elevation (array of 96 × 16 LEDs, emission peak: 470 nm). At the display horizontal midline, each LED subtended an angle of 3.75° on the fly’s retina. The fly was suspended between two magnets that maintained the animal in place and free to rotate about its vertical axis (i.e., yaw axis). The pin was attracted at its top end toward the upper magnetic pivot and its moment of inertia encompassed less than 1% of the fly’s moment of inertia (Bender and Dickinson, 2006; Fry et al., 2003). The fly was illuminated from below with an array of eight infrared diodes (emission peak: 940 nm) and recorded from the bottom with an infrared-sensitive camera (Blackfly S USB3, Teledyne FLIR) fitted with a zoom lens (InfiniStix 0.5x/0.024, Infinity Photo-Optical) at 200 frames s^-1^. The lens also held a long pass filter to block the light emitted by the display (NIR, Edmund Optics). At the beginning of each trial, after ten seconds of acclimatation, the fly was presented for 20 s with a rotating wide-field panorama, which elicited strong optomotor responses, in each direction (CW and CCW). Flies whose behavior was characterized by excessive wobble indicating poor tethering, were discarded.

### Magnetic-tether visual stimuli

Visual stimuli consisted in a motion-defined bar (30° x 56°) and a wide-field panorama (360° x 56°) of random bright and dark vertical stripes rotating horizontally around the fly in CW and CCW directions at 112.5° s^-1^. We decided to use these stimuli because previous experiments conducted in our laboratory showed that motion-defined bars elicit a higher number of body-saccades compared to solid luminance-defined bars (∼ 50% more) (Mongeau and Frye, 2017). For similar reasons, the speed selected elicits robust bar tracking behavior (Mongeau and Frye, 2017). Both stimuli were generated, as mentioned above, using custom-written MATLAB code and randomly chosen during each trial among 96 different patterns. Each trial involved 25 s of a rotating stimulus at constant speed and 5 s of resting period in which the stimulus stopped moving. For the bar, the initial position was selected from a pseudo-random sequence. Each fly was tested in four different and randomized trials (2 stimuli x 2 directions) without repetitions to minimize habituation. The full experiment lasted ∼ 3 min. Only flies that either flew continuously or stopped briefly only once were included in the analysis.

### Optogenetic stimulation

Optogenetic experiments were conducted in a setup similar to the one used for magnetic-tether experiments. Likewise, a cylindrical LED panel display (array of 96 × 16 LEDs, emission peak: 470 nm) was used and two layers of neutral filter were placed over the display to reduce the light intensity. A red LED (emission peak: 685 nm, 4V) was positioned laterally to the post bearing the magnetic pivot within which the fly was suspended. Its beam covered the entire fly and, since the fly freely rotated around the yaw axis. Thus, the angle of illumination varied during flight. We used the same visual stimuli, speed, directions and trial duration used in magnetic-tether experiments. On top of visual rotating stimuli, flies were exposed to contiguous epochs of LED On and Off which lasted 5 s each. The starting optogenetic epoch was randomly selected so that flies could start the trial with the LED On or Off. This means that during the 25 s of visual stimulation, flies could be optogenetically stimulated for either 10 s or 15 s in total. After every stimulation, flies were left to rest for 5 s facing a static random-stripes pattern. Moreover, we included trials where the LED remained Off throughout the visual stimulation. The experiment lasted ∼ 6 min (30 s x 3 LED intensities x 2 stimuli x 2 directions). As done for the magnetic-tether experiments, we discarded flies that showed signs of poor tethering preparation (i.e., asymmetric wing beat amplitude). All-trans-retinal (ATR) is required to get a proper CsChrimson protein conformation. In order to boost flies’ performance, although flies endogenously produce retinal, we added ATR to the food. The progeny from the crosses between driver lines and UAS-CsChrimson reporter were raised in the darkness to avoid the channel’s stimulation. After eclosion, newborn flies were transferred in 0.5 mM ATR food and kept there for 3-5 days until the experiment.

### Behavioral analysis

Data collected either in the rigid-tether or in the magno-tether setups were analyzed in RStudio (RStudio Team, 2021) using custom R scripts. Axon binary files (.abf) from the DAQ used in rigid tether experiments were imported by using the R package *abf2* (Caldwell, 2015) and pre-processed (data filtering) as previously described (Keleş et al., 2018). Video recordings from the camera on the magno-tether setup were imported into MATLAB and the flies’ heading offline tracked by using custom scripts. Video heading files and DAQ files were then imported into RStudio by using the R package *R*.*matlab* (Bengtsson, 2018). Saccade detection and tracking bouts were analyzed as previously described (Mongeau and Frye, 2017). Data plotting was performed by R package *ggplot2* (Wickham, 2016).

### Model

In the integrator-and-fire model physiologically-inspired on T3 neurons, the nonlinear least-squares estimates of the parameters of the fitted calcium imaging responses were computed in RStudio (RStudio Team, 2021) using a nonlinear regression analysis (Bates, 1988). We used the *nls* function with a predefined model formula inspired to the probability density function of a Beta distribution *y* = (*x*^(*α* − 1)^ × (1 − *x*)(^*β* − 1)^) × *γ*, where *α*, and *β* are shape parameters, while *γ* is a scaling factor. These models minimized the number of parameters, while maintaining wide flexibility and goodness of fit. The behavioral control model was simulated in Simulink (MATLAB).

### Statistics

Generalized linear mixed effects (GLME) models were used to fit the data in order to consider the random effects represented by individual flies. GLME models avoid averaging that reduces the statistical power and allows for adjusting the estimates for repeated sampling and for sample imbalancing. We fitted the data using the R package *lme4* (Bates et al., 2015). Analysis of Variance (ANOVA) was computed on the estimated parameters by the models. Pairwise post-hoc comparisons corrected with Bonferroni method on the fixed effects of the models were then performed using the R package *emmeans* (Lenth, 2021). In violin-box plots, mean and median were reported as central tendency measures, bottom and top edges of the box represent 25th (Q1) and 75th (Q3) percentiles and whiskers represent the lowest and highest datum within 1.5 interquartile range (Q3 - Q1).

**Supplementary Table 1.** Origin of reagents used in this study.

| Reagent type | Source | Identifier |
| --- | --- | --- |
| *D. melanogaster*: Empty-SplitGal4 [(p65ADZp.Uw(attP40); ZpGDBD.Uw(attP2))] | Bloomington<br>Drosophila<br>Stock Center | BDSC<br>#79603 |
| *D. melanogaster*: T3-SplitGal4 [VT002055-p65ADZp(attP40); R65B04- ZpGDBD(attP2)] | (Keleş et al., 2020) | N/A |
| *D. melanogaster*: T2a-SplitGal4 [VT012791-p65ADZp(attP40); R47E02- ZpGDBD(attP2)] | (Keleş et al., 2020) | N/A |
| *D. melanogaster*: T4/T5-SplitGal4 [R59E08-p65.AD(attP40); R42F06-Gal4.DBD(attP2)] | G. Rubin | N/A |
| *D. melanogaster*: 20XUAS-IVS-GCaMP6f(VK00005) | Bloomington<br>Drosophila<br>Stock Center | BDSC<br>#52869 |
| *D. melanogaster*: 10XUAS-IVS-eGFP-Kir2.1(attP2) | (von Reyn et al., 2017) | N/A |
| *D. melanogaster*: 20xUAS-Chrimson::tdTomato(VK00005) | D. Anderson | N/A |

**Supplementary Table 2.**
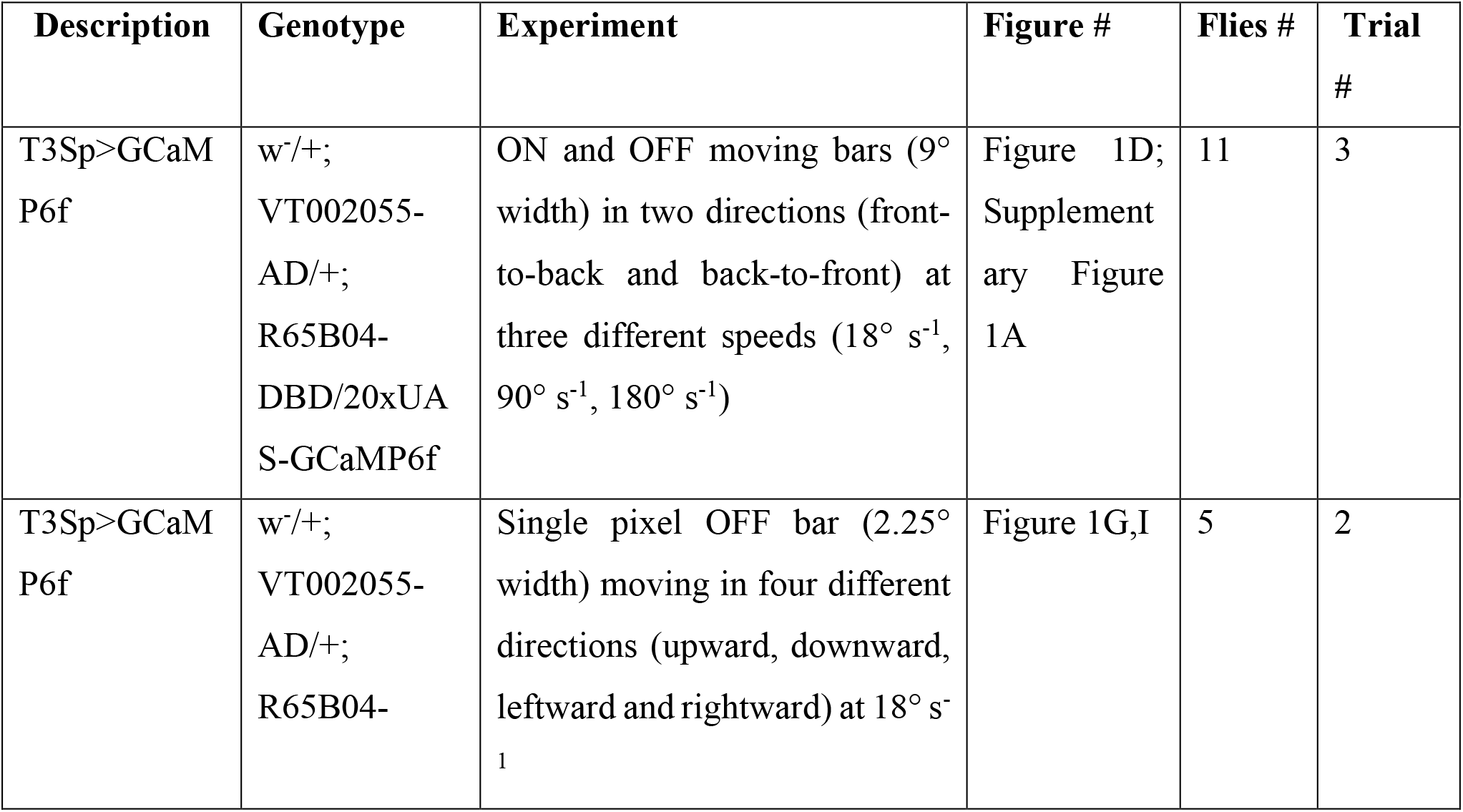
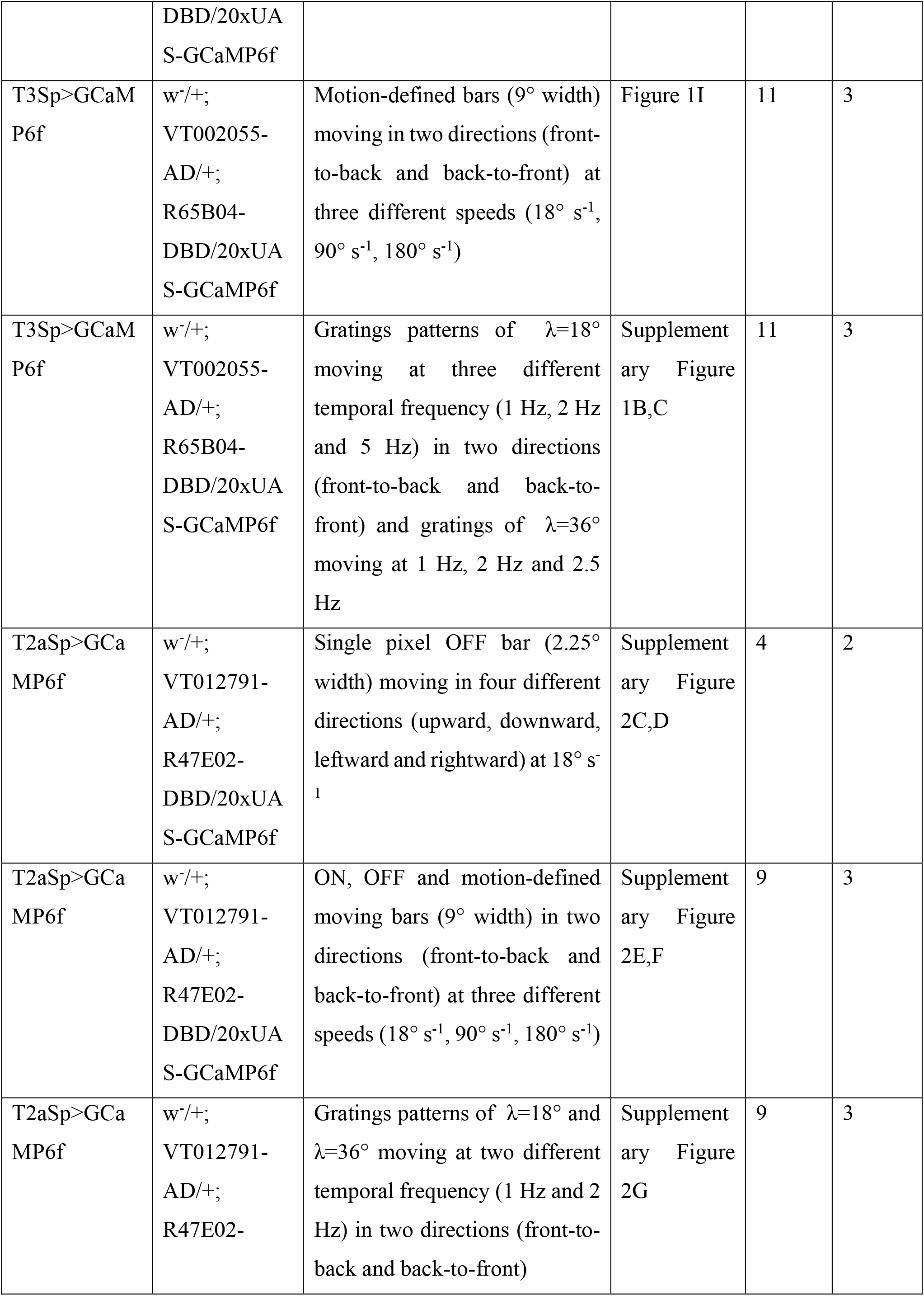
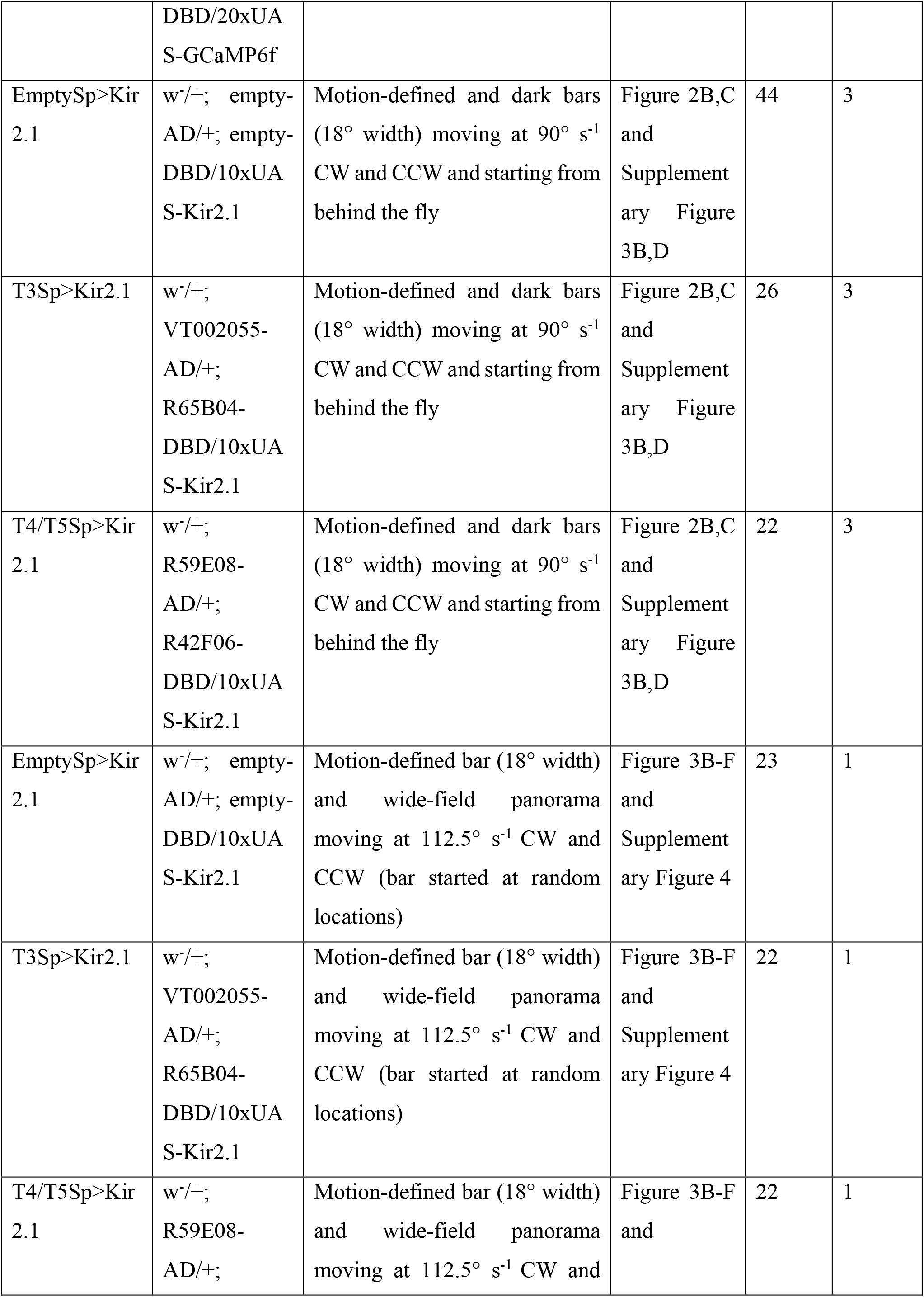
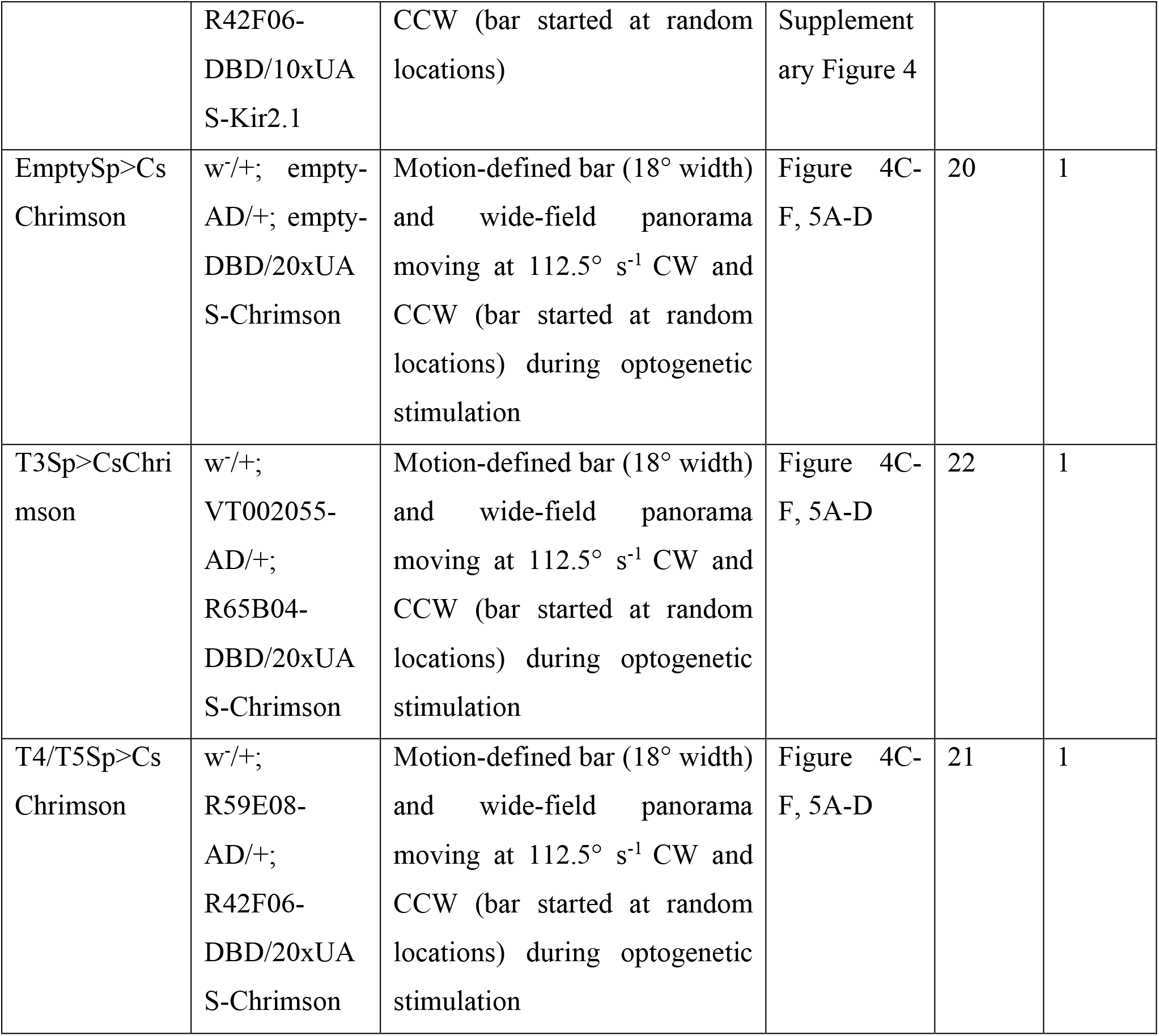
Genotypes and experimental parameters used in each figure.

## Acknowledgements

We thank Ivan Lopez for preliminary experimental results, Mehmet F. Keleş for reagents, and Ben J. Hardcastle for technical advice. This work was supported by a grant from the National Institutes of Health (R01-EY026031) to M.A.F.

## Author contributions

G.F., and M.A.F. conceived the project. G.F. performed two-photon imaging, behavioral experiments in rigidly and magnetically tethered flies, and analyzed the data. G.F. and M.A.F. interpreted experiments and wrote the paper.

## Videos

**Video 1**. Calcium imaging recording from T3 dendrites. T3 neurons expressing GCaMP6f respond to a back-to-front moving bright bar at 90° s^-1^.

**Video 2**. EmpySp>Kir2.1 fly presenting with a revolving motion-defined bar. Left: bottom view of a single fly within an animated cartoon of the surrounding display presenting a motion-defined bar revolving for 25 s at 112.5° s^-1^. Right-top: fly heading (white) and bar position (gray). Right-bottom: Error (red) between bar position and fly heading. EmptySp flies track the bar by using a saccade-and-fixation strategy. Original video recorded at 200 fps.

**Video 3**. T3Sp>Kir2.1 fly presenting with a revolving motion-defined bar. Conditions identical to Supplementary Video 2. T3Sp flies do not track the bar.

**Video 4**. T4/T5Sp>Kir2.1 fly presenting with a revolving motion-defined bar. Conditions identical to Supplementary Video 2. T4/T5Sp flies track the bar with some defects in gaze stabilization.

## Data availability

Data will be posted in an online repository upon publication.

## Code availability

Analysis and plotting code will be posted in an online repository upon publication.

---

**Supplementary Figure 1.**
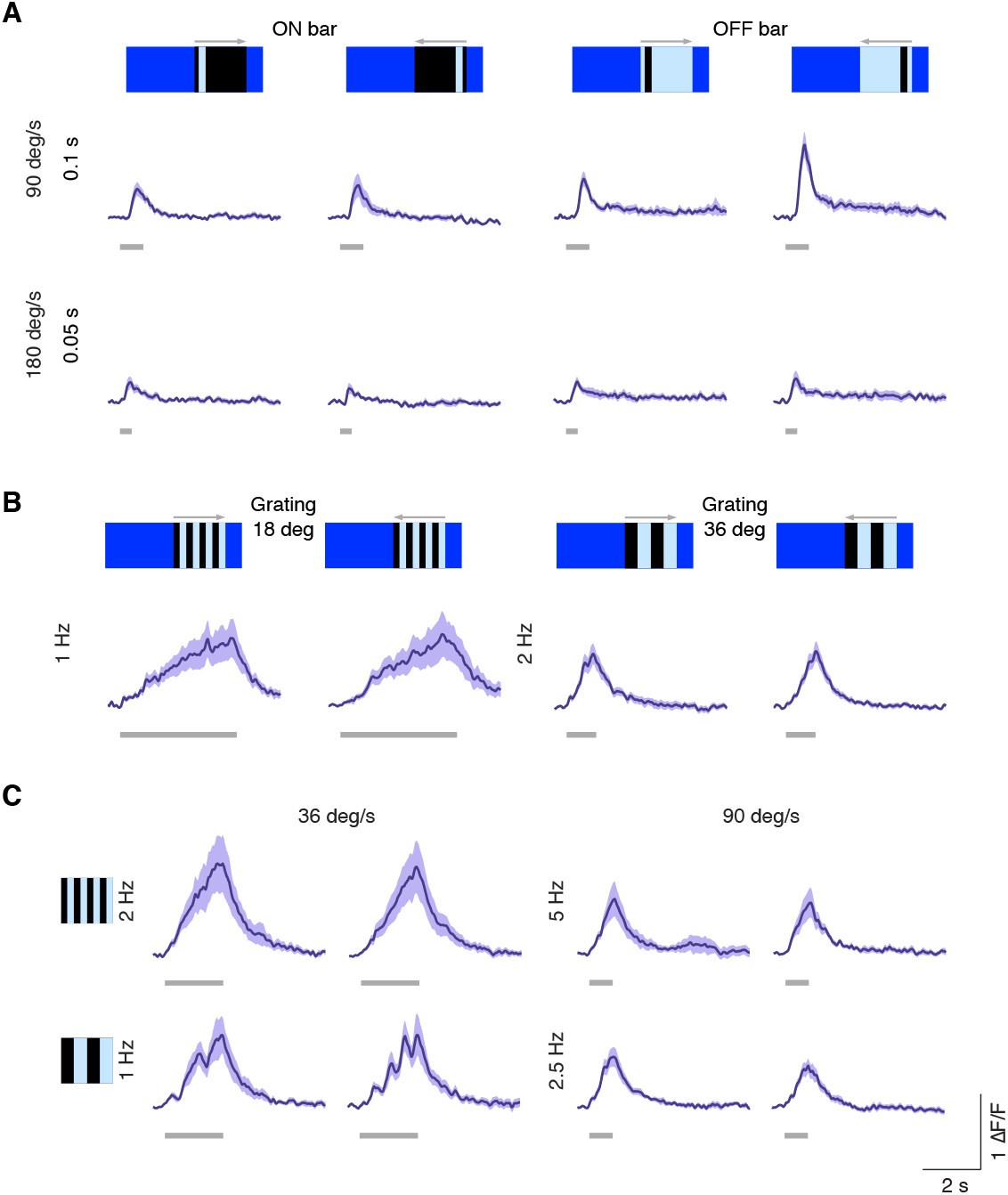
Speed tuning of T3 neurons. (**A**) Average responses (mean ± s.e.m.) of T3 neurons to ON and OFF solid moving bars (9° x 72°, width x height) moving in two different directions (front-to-back and back-to-front) at two different speeds. Visual stimuli are depicted at the top. Light gray horizontal bars at the bottom indicate stimulus presentation (n = 11 flies, 3 repetitions per fly). (**B**) T3 responses to moving gratings of different spatial and temporal frequencies. (**C**) Left: T3 neurons show similar peaks for gratings moving at 36° s^-1^ regardless of the spatial frequency of the stimuli. Right: same effect for gratings moving at 90° s^-1^. The slow calcium integration dynamic combined with the full rectification represents an optimal mechanism for speed detection.

**Supplementary Figure 2.**
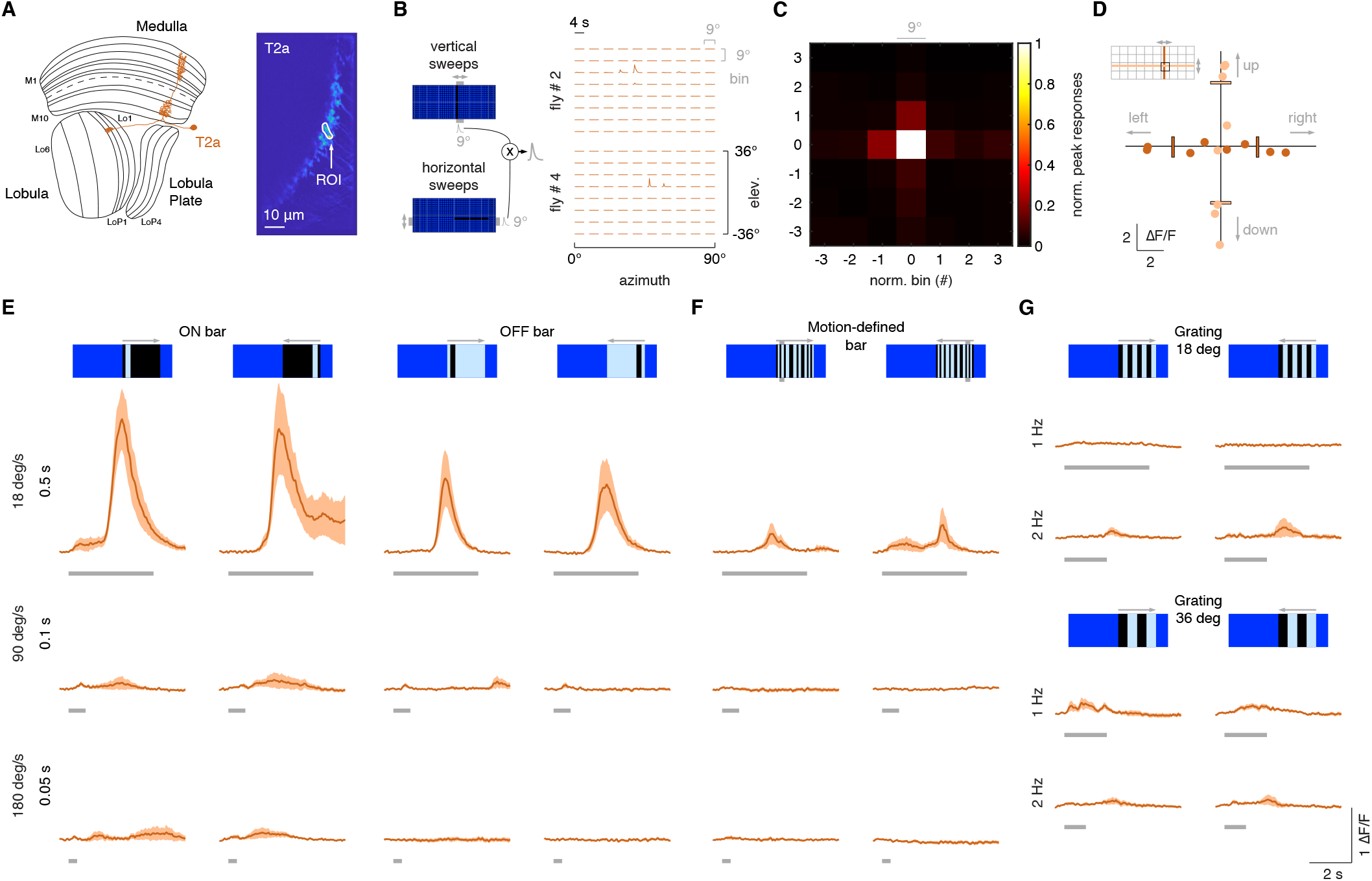
T2a neurons do not show a broad temporal sensitivity. (**A**) Left: schematic representation of a T2a neuron (orange) within the optic lobe. Right: ROI drawn around the presynaptic terminal in the lobula of a T2a neuron expressing GCaMP6f. Image representing the mean activity from the two-photon imaging experiment in a representative fly. (**B**) Left: representation of the procedure used to probe the RF of T2a (as done in **Figure 1F**). Right: matrix of the responses obtained by multiplying horizontal and vertical sweeps in two representative flies. (**C**) Mean of the normalized peak responses of T2a neurons by spatial location (n = 4 flies). Bin = 0 represents the center of the RF. (**D**) Directional calcium peak responses to a 2.25° dark bar moving (18° s^-1^) in the four cardinal directions of individual flies. (**E**) Average responses (mean ± s.e.m.) to moving ON and OFF solid bars (9° x 72°, width x height) at different speeds in two different directions (front-to-back and back-to-front). Visual stimuli are depicted at the top. Light gray horizontal bars at the bottom indicate stimulus presentation (n = 9 flies, 3 repetitions per fly). (**F**) Average responses (mean ± s.e.m.) to motion-defined bars moving in two different directions (front-to-back and back-to-front) at different speeds. (**G**) Top: T2a responses to a grating of λ=18° moving front-to-back and back-to-front at two different temporal frequencies. Bottom: T2a responses to a grating of λ=36°.

**Supplementary Figure 3.**
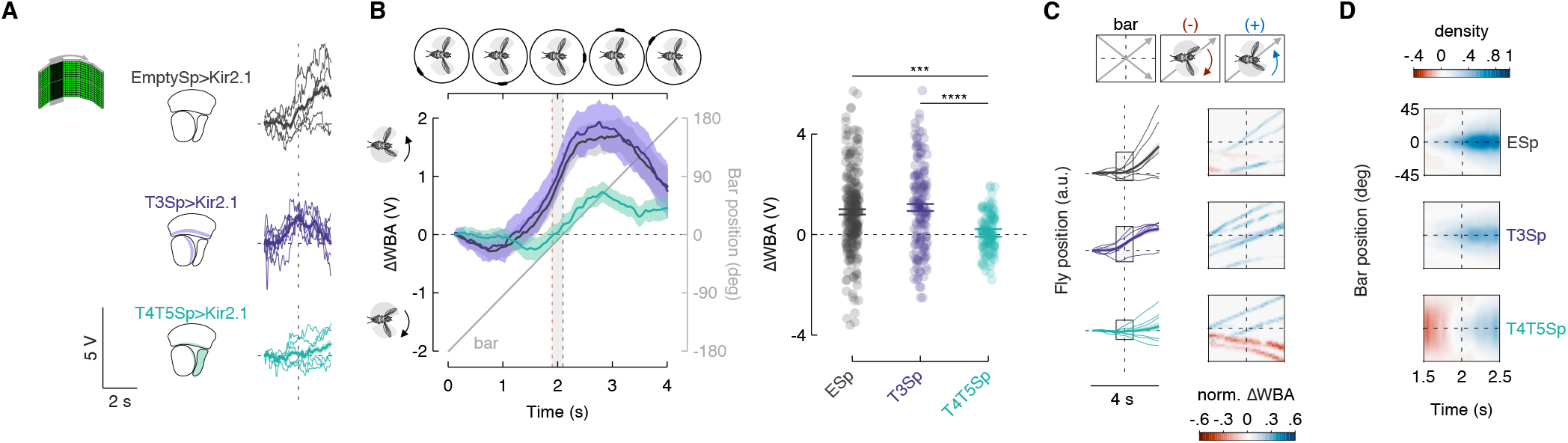
Hyperpolarization of T3 does not compromise the syn-directional response. (**A**) Left: representation of the visual stimulus (dark bar revolving at 90° s^-1^). Middle: schematic representation of the optic lobe regions where Kir2.1 channels were expressed in the three genotypes tested. Right: single trials (3 repetitions x 2 directions) of ΔWBA responses (thin lines) to rotation of a luminance-defined bar in three representative flies (T4/T5Sp>Kir2.1 data are reproduced from Keleş et al., 2018). Thick line represents the mean (responses to CCW rotations were reflected and pooled with CW responses). Vertical dashed lines indicate when the bar is at the fly’s visual midline while the horizontal ones represent ΔWBA = 0. (**B**) Left: population average time series steering responses (mean ± s.e.m.) in the three genotypes tested (T4/T5Sp data replotted from Keleş et al., 2018) to a luminance-defined bar. Gray shaded region (between vertical red and black dashed lines) represents a 200 ms time window before the bar crosses the fly’s visual midline (n = 44 EmptySp>Kir2.1, n = 26 T3Sp>Kir2.1, n = 22 T4/T5Sp>Kir2.1). T4/T5Sp>Kir2.1 reduces syn-directional anticipatory steering, whereas T3Sp>Kir2.1 shows normal syn-directional steering. Right: Dot plot average ΔWBA values across the 200 ms time window per trial. Dark dots indicate the mean and the horizontal bars indicate s.e.m. (F_(2, 89)_ = 14.48, p < .0001; EmptySp vs T3Sp: p = .94; EmptySp vs T4/T5Sp: p < .0001; T3Sp vs T4/T5Sp: p < .0001). (**C**) Left: arbitrary fly position (thin lines) resulting from the integration of ΔWBA values over time in three representative flies. Thick lines represent the mean. Right: space-time plot of the normalized fly position within the gray shaded boxes highlighted to the left. Color-code represents the direction of the steering effort (red: counter-directional; blue: syn-directional). (**D**) Heat maps of flies’ steering effort at the population level in the three genotypes as a function of the bar position. EmptySp and T3Sp flies show a strong syn-directional response (blue blob) while T4/T5Sp flies show an early counter-directional response and a weak syn-directional response.

**Supplementary Figure 4.**
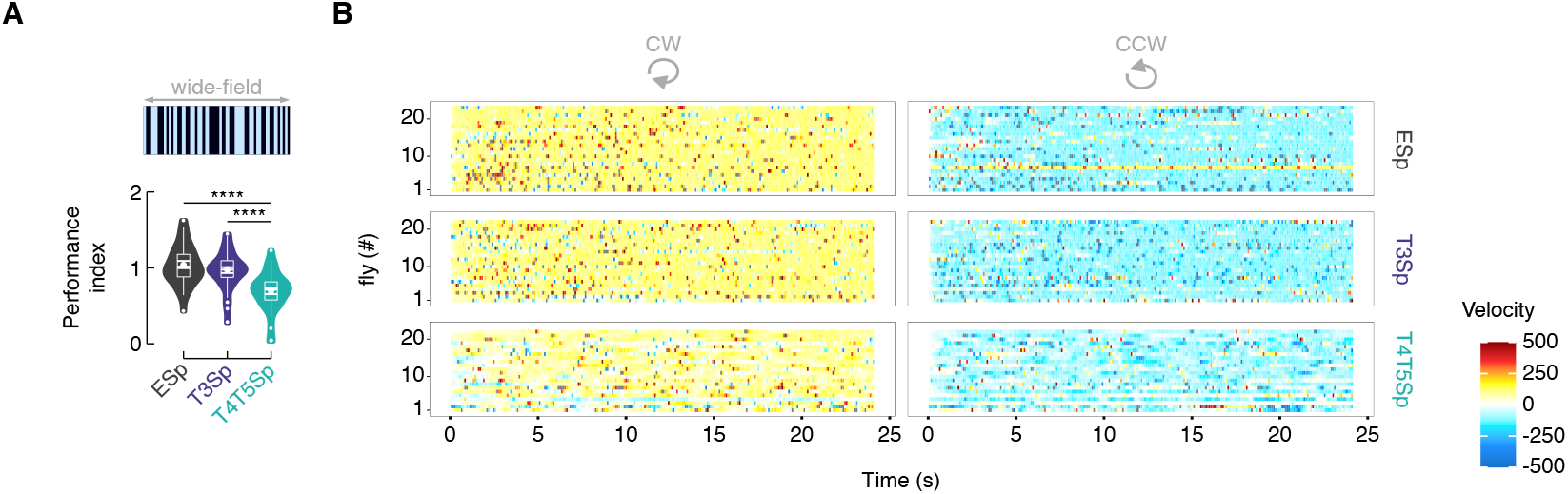
T3 silencing does not affect the response to the rotation of a wide-field panorama. (**A**) Violin-box plot of the performance index (i.e., gain) to a rotating random wide-field pattern of dark and bright stripes in the three genotypes. T4/T5Sp shows a strong reduction of the smooth tracking gain while T3Sp shows a normal tracking response (pairwise post-hoc comparisons adjusted Bonferroni, EmptySp vs T3Sp: p = 1; EmptySp vs T4/T5Sp: p < .0001; T3Sp vs T4/T5Sp: p < .0001). (**B**) Raster plot of the velocity (yellow-red: CW; green-blue: CCW) per fly during the rotation of the wide-field panorama (each bin represents 100 ms of average velocity). Note that T4/T5Sp>Kir2.1 flies reduce the tracking velocity, whereas T3Sp>Kir2.1 flies show a velocity comparable to control flies (n = 20 EmptySp>Kir2.1, n = 22 T3Sp>Kir2.1, n = 21 T4/T5>Kir2.1).

**Supplementary Figure 5.**
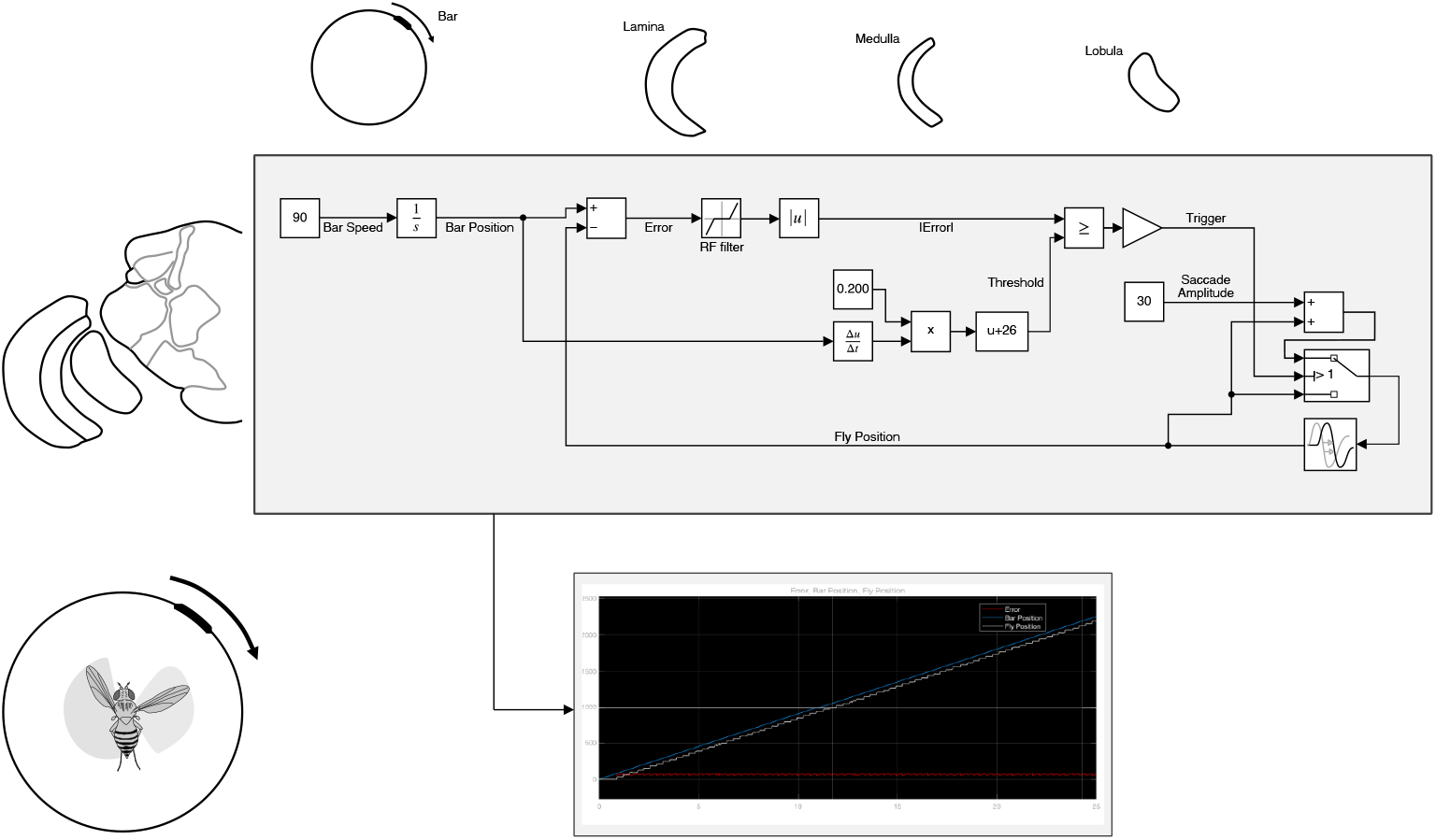
Control model for triggering saccades. Top: simulink (MATLAB) implementation of the physiologically-inspired model in **Figure 6**. The bar speed (90° s^-1^) is integrated over a quite narrow time window (∼ 200 ms). This means that T3 neurons might encode the bar speed and a downstream partner might integrate this information over a selective amount of time, encoding the bar position and triggering a saccade when the amount of calcium reaches a specific threshold. In this model saccade amplitudes are considered a fixed parameter (30°) but in an alternative version they could be easily tuned to the bar speed (as a role played by the T4/T5 pathway). Bottom: simulation of the fly behavior (angular position) in a bar tracking task according to the control model.

## References

Agrochao M, Tanaka R, Salazar-Gatzimas E, Clark DA. 2020. Mechanism for analogous illusory motion perception in flies and humans. Proc Natl Acad Sci U S A 117:23044–23053.

Aptekar JW, Frye MA. 2013. Higher-order figure discrimination in fly and human vision. Curr Biol 23:R694–700.

Aptekar JW, Keleş MF, Lu PM, Zolotova NM, Frye MA. 2015. Neurons forming optic glomeruli compute figure-ground discriminations in Drosophila. J Neurosci 35:7587–7599.

Aptekar JW, Shoemaker PA, Frye MA. 2012. Figure tracking by flies is supported by parallel visual streams. Curr Biol 22:482–487.

Bahl A, Ammer G, Schilling T, Borst A. 2013. Object tracking in motion-blind flies. Nat Neurosci 16:730–738.

Bahl A, Serbe E, Meier M, Ammer G, Borst A. 2015. Neural Mechanisms for Drosophila Contrast Vision. Neuron 88:1240–1252.

Baines RA, Uhler JP, Thompson A, Sweeney ST, Bate M. 2001. Altered electrical properties in Drosophila neurons developing without synaptic transmission. J Neurosci 21:1523–1531.

Bates DM. 1988. Nonlinear regression analysis and its applications. New York: Wiley.

Bates D, Mächler M, Bolker B, Walker S. 2015. Fitting Linear Mixed-Effects Models Using lme4. J Stat Softw 67:1–48.

Bender JA, Dickinson MH. 2006. Visual stimulation of saccades in magnetically tethered Drosophila. J Exp Biol 209:3170–3182.

Bengtsson H. 2018. R.matlab: Read and Write MAT Files and Call MATLAB from Within R. Comprehensive R Archive Network (CRAN).

Borst A. 2014. Fly visual course control: behaviour, algorithms and circuits. Nat Rev Neurosci 15:590–599.

Borst A, Drews M, Meier M. 2020a. The neural network behind the eyes of a fly. Current Opinion in Physiology 16:33–42.

Borst A, Haag J, Mauss AS. 2020b. How fly neurons compute the direction of visual motion. J Comp Physiol A Neuroethol Sens Neural Behav Physiol 206:109–124.

Busch C, Borst A, Mauss AS. 2018. Bi-directional Control of Walking Behavior by Horizontal Optic Flow Sensors. Curr Biol 28:4037–4045.e5.

Caldwell M. 2015. abf2: Load Gap-Free Axon ABF2 Files.

Cheong HS, Siwanowicz I, Card GM. 2020. Multi-regional circuits underlying visually guided decision-making in Drosophila. Curr Opin Neurobiol 65:77–87.

Demerec M. 2008. Biology of Drosophila. Cold Spring Harbor, NY, USA: Cold Spring Harbor Laboratory Press.

Duistermars BJ, Frye M. 2008. A magnetic tether system to investigate visual and olfactory mediated flight control in Drosophila. J Vis Exp. doi:10.3791/1063

Egelhaaf M. 1985. On the neuronal basis of figure-ground discrimination by relative motion in the visual system of the fly. Biol Cybern 52:195–209.

Fenk LM, Poehlmann A, Straw AD. 2014. Asymmetric processing of visual motion for simultaneous object and background responses. Curr Biol 24:2913–2919.

Ferreira CH, Moita MA. 2020. Behavioral and neuronal underpinnings of safety in numbers in fruit flies. Nat Commun 11:4182.

Fischbach K-F, Dittrich APM. 1989. The optic lobe of Drosophila melanogaster. I. A Golgi analysis of wild-type structure. Cell Tissue Res 258:441–475.

Fisher YE, Silies M, Clandinin TR. 2015. Orientation Selectivity Sharpens Motion Detection in Drosophila. Neuron 88:390–402.

Fry SN, Sayaman R, Dickinson MH. 2003. The aerodynamics of free-flight maneuvers in Drosophila. Science 300:495–498.

Geurten BRH, Jähde P, Corthals K, Göpfert MC. 2014. Saccadic body turns in walking Drosophila. Front Behav Neurosci 8:365.

Groschner LN, Malis JG, Zuidinga B, Borst A. 2022. A biophysical account of multiplication by a single neuron. Nature. doi:10.1038/s41586-022-04428-3

Gruntman E, Romani S, Reiser MB. 2019. The computation of directional selectivity in the Drosophila OFF motion pathway. Elife 8. doi:10.7554/eLife.50706

Guizar-Sicairos M, Thurman ST, Fienup JR. 2008. Efficient subpixel image registration algorithms. Opt Lett 33:156–158.

Haikala V, Joesch M, Borst A, Mauss AS. 2013. Optogenetic control of fly optomotor responses. J Neurosci 33:13927–13934.

Heisenberg M, Wonneberger R, Wolf R. 1978. Optomotor-blindH31—a Drosophila mutant of the lobula plate giant neurons. J Comp Physiol 124:287–296.

Joesch M, Schnell B, Raghu SV, Reiff DF, Borst A. 2010. ON and OFF pathways in Drosophila motion vision. Nature 468:300–304.

Keleş MF, Frye MA. 2017a. The eyes have it. Elife. doi:10.7554/eLife.24896

Keleş Mf, Frye MA. 2017b. Object-Detecting Neurons in Drosophila. Curr Biol 27:680–687.

Keleş MF, Hardcastle BJ, Städele C, Xiao Q, Frye MA. 2020. Inhibitory Interactions and Columnar Inputs to an Object Motion Detector in Drosophila. Cell Rep 30:2115–2124.e5.

Keleş MF, Mongeau J-M, Frye MA. 2018. Object features and T4/T5 motion detectors modulate the dynamics of bar tracking by Drosophila. J Exp Biol. doi:10.1242/jeb.190017

Klapoetke NC, Murata Y, Kim SS, Pulver SR, Birdsey-Benson A, Cho YK, Morimoto TK, Chuong AS, Carpenter EJ, Tian Z, Wang J, Xie Y, Yan Z, Zhang Y, Chow By, Surek B, Melkonian M, Jayaraman V, Constantine-Paton M, Wong GK-S, Boyden ES. 2014. Independent optical excitation of distinct neural populations. Nat Methods 11:338–346.

Klapoetke NC, Nern A, Rogers EM, Rubin GM, Reiser MB, Card GM. 2022. A functionally ordered visual feature map in the Drosophila brain. Neuron S0896–6273(22)00178–7.

Konstantinides N, Kapuralin K, Fadil C, Barboza L, Satija R, Desplan C. 2018. Phenotypic Convergence: Distinct Transcription Factors Regulate Common Terminal Features. Cell 174:622–635.e13.

Land MF. 1992. Visual tracking and pursuit: Humans and arthropods compared. J Insect Physiol 38:939–951.

Lenth RV. 2021. emmeans: Estimated Marginal Means, aka Least-Squares Means.

Liang P, Heitwerth J, Kern R, Kurtz R, Egelhaaf M. 2012. Object representation and distance encoding in three-dimensional environments by a neural circuit in the visual system of the blowfly. J Neurophysiol 107:3446–3457.

Maisak MS, Haag J, Ammer G, Serbe E, Meier M, Leonhardt A, Schilling T, Bahl A, Rubin GM, Nern A, Dickson BJ, Reiff DF, Hopp E, Borst A. 2013. A directional tuning map of Drosophila elementary motion detectors. Nature 500:212–216.

Mauss AS, Borst A. 2020. Optic flow-based course control in insects. Curr Opin Neurobiol 60:21–27.

Mongeau J-M, Cheng KY, Aptekar J, Frye MA. 2019. Visuomotor strategies for object approach and aversion in Drosophila melanogaster. J Exp Biol 222. doi:10.1242/jeb.193730

Mongeau J-M, Frye MA. 2017. Drosophila Spatiotemporally Integrates Visual Signals to Control Saccades. Curr Biol 27:2901–2914.e2.

Panser K, Tirian L, Schulze F, Villalba S, Jefferis GSXE, Bühler K, Straw AD. 2016. Automatic Segmentation of Drosophila Neural Compartments Using GAL4 Expression Data Reveals Novel Visual Pathways. Curr Biol 26:1943–1954.

Reichardt W, Poggio T. 1976. Visual control of orientation behaviour in the fly. Part I. A quantitative analysis. Q Rev Biophys 9:311–75, 428–38.

Reichardt W, Poggio T, Hausen K. 1983. Figure-ground discrimination by relative movement in the visual system of the fly. Biol Cybern 46:1–30.

Reiser MB, Dickinson MH. 2010. Drosophila fly straight by fixating objects in the face of expanding optic flow. J Exp Biol 213:1771–1781.

Reiser MB, Dickinson MH. 2008. A modular display system for insect behavioral neuroscience. J Neurosci Methods 167:127–139.

RStudio Team. 2021. RStudio: Integrated Development Environment for R. Boston, MA: RStudio, PBC.

Shinomiya K, Huang G, Lu Z, Parag T, Xu CS, Aniceto R, Ansari N, Cheatham N, Lauchie S, Neace E, Ogundeyi O, Ordish C, Peel D, Shinomiya A, Smith C, Takemura S, Talebi I, Rivlin PK, Nern A, Scheffer LK, Plaza SM, Meinertzhagen IA. 2019. Comparisons between the ON-and OFF-edge motion pathways in the Drosophila brain. Elife 8. doi:10.7554/eLife.40025

Shinomiya K, Nern A, Meinertzhagen IA, Plaza SM, Reiser MB. 2022. Neuronal circuits integrating visual motion information in Drosophila melanogaster. Current Biology. doi:10.1016/j.cub.2022.06.061

Städele C, Keleş MF, Mongeau J-M, Frye MA. 2020. Non-canonical Receptive Field Properties and Neuromodulation of Feature-Detecting Neurons in Flies. Curr Biol. doi:10.1016/j.cub.2020.04.069

Strother JA, Wu S-T, Wong AM, Nern A, Rogers EM, L. JQ, Rubin GM, Reiser MB. 2017. The Emergence of Directional Selectivity in the Visual Motion Pathway of Drosophila. Neuron 94:168–182.e10.

Takemura S-Y, Bharioke A, Lu Z, Nern A, Vitaladevuni S, Rivlin PK, Katz WT, Olbris DJ, Plaza SM, Winston P, Zhao T, Horne JA, Fetter RD, Takemura S, Blazek K, Chang L-A, Ogundeyi O, Saunders MA, Shapiro V, Sigmund C, Rubin GM, Scheffer LK, Meinertzhagen IA, Chklovskii DB. 2013. A visual motion detection circuit suggested by Drosophila connectomics. Nature 500:175–181.

Takemura S-Y, Xu CS, Lu Z, Rivlin PK, Parag T, Olbris DJ, Plaza S, Zhao T, Katz WT, Umayam L, Weaver C, Hess HF, Horne JA, Nunez-Iglesias J, Aniceto R, Chang L-A, Lauchie S, Nasca A, Ogundeyi O, Sigmund C, Takemura S, Tran J, Langille C, Le Lacheur K, McLin S, Shinomiya A, Chklovskii DB, Meinertzhagen IA, Scheffer LK. 2015. Synaptic circuits and their variations within different columns in the visual system of Drosophila. Proc Natl Acad Sci U S A 112:13711–13716.

Tammero LF, Dickinson MH. 2002. The influence of visual landscape on the free flight behavior of the fruit fly Drosophila melanogaster. J Exp Biol 205:327–343.

Tammero LF, Frye MA, Dickinson MH. 2004. Spatial organization of visuomotor reflexes in Drosophila. J Exp Biol 207:113–122.

Tanaka R, Clark DA. 2022. Neural mechanisms to exploit positional geometry for collision avoidance. Curr Biol 32:2357–2374.e6.

Tanaka R, Clark DA. 2020. Object-Displacement-Sensitive Visual Neurons Drive Freezing in Drosophila. Curr Biol. doi:10.1016/j.cub.2020.04.068

Theobald JC, Duistermars BJ, Ringach DL, Frye MA. 2008. Flies see second-order motion. Curr Biol 18:R464–5.

van Breugel F, Dickinson MH. 2012. The visual control of landing and obstacle avoidance in the fruit fly Drosophila melanogaster. J Exp Biol 215:1783–1798.

von Reyn CR, Nern A, Williamson WR, Breads P, Wu M, Namiki S, Card GM. 2017. Feature Integration Drives Probabilistic Behavior in the Drosophila Escape Response. Neuron 94:1190–1204.e6.

Weir PT, Dickinson MH. 2015. Functional divisions for visual processing in the central brain of flying Drosophila. Proc Natl Acad Sci U S A 112:E5523–5532.

Wickham H. 2016. ggplot2: Elegant Graphics for Data Analysis. Springer-Verlag New York.

Wu M, Nern A, Williamson WR, Morimoto MM, Reiser MB, Card GM, Rubin GM. 2016. Visual projection neurons in the Drosophila lobula link feature detection to distinct behavioral programs. Elife 5. doi:10.7554/eLife.21022

Yang HH, Clandinin TR. 2018. Elementary Motion Detection in Drosophila: Algorithms and Mechanisms. Annu Rev Vis Sci 4:143–163.

