## Supplemental figures for "Feature detecting columnar neurons mediate object tracking saccades in *Drosophila*"

### Supplementary Figure 1

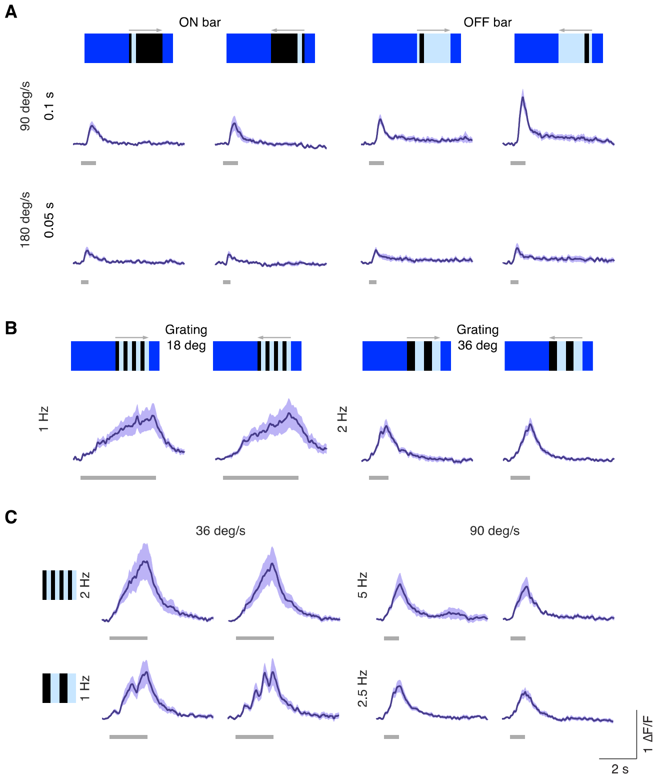


**Supplementary Figure 1.** Speed tuning of T3 neurons. (**A**) Average responses (mean ± s.e.m.) of T3 neurons to ON and OFF solid moving bars (9° x 72°, width x height) moving in two different directions (front-to-back and back-to-front) at two different speeds. Visual stimuli are depicted at the top. Light gray horizontal bars at the bottom indicate stimulus presentation (n = 11 flies, 3 repetitions per fly). (**B**) T3 responses to moving gratings of different spatial and temporal frequencies. (**C**) Left: T3 neurons show similar peaks for gratings moving at 36° $s^{-1}$ regardless of the spatial frequency of the stimuli. Right: same effect for gratings moving at 90° $s^{-1}$. The slow calcium integration dynamic combined with the full rectification represents an optimal mechanism for speed detection.

### Supplementary Figure 2

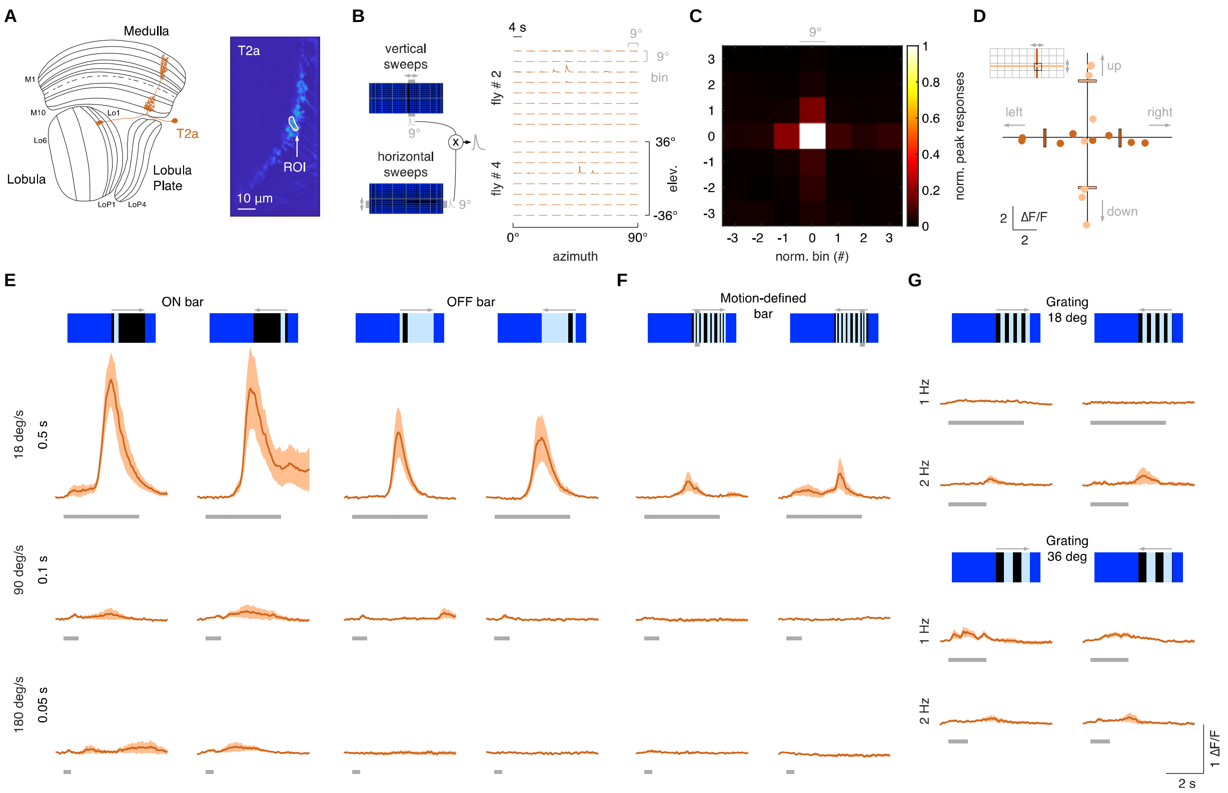


**Supplementary Figure 2.** T2a neurons do not show a broad temporal sensitivity. (**A**) Left: schematic representation of a T2a neuron (orange) within the optic lobe. Right: ROI drawn around the presynaptic terminal in the lobula of a T2a neuron expressing GCaMP6f. Image representing the mean activity from the two-photon imaging experiment in a representative fly. (**B**) Left: representation of the procedure used to probe the RF of T2a (as done in **Figure 1F**). Right: matrix of the responses obtained by multiplying horizontal and vertical sweeps in two representative flies. (**C**) Mean of the normalized peak responses of T2a neurons by spatial location (n = 4 flies). Bin = 0 represents the center of the RF. (**D**) Directional calcium peak responses to a 2.25° dark bar moving (18° $s^{-1}$) in the four cardinal directions of individual flies. (**E**) Average responses (mean ± s.e.m.) to moving ON and OFF solid bars (9° x 72°, width x height) at different speeds in two different directions (front-to-back and back-to-front). Visual stimuli are depicted at the top. Light gray horizontal bars at the bottom indicate stimulus presentation (n = 9 flies, 3 repetitions per fly). (**F**) Average responses (mean ± s.e.m.) to motion-defined bars moving in two different directions (front-to-back and back-to-front) at different speeds. (**G**) Top: T2a responses to a grating of $\lambda$=18° moving front-to-back and back-to-front at two different temporal frequencies. Bottom: T2a responses to a grating of $\lambda$=36°.

### Supplementary Figure 3

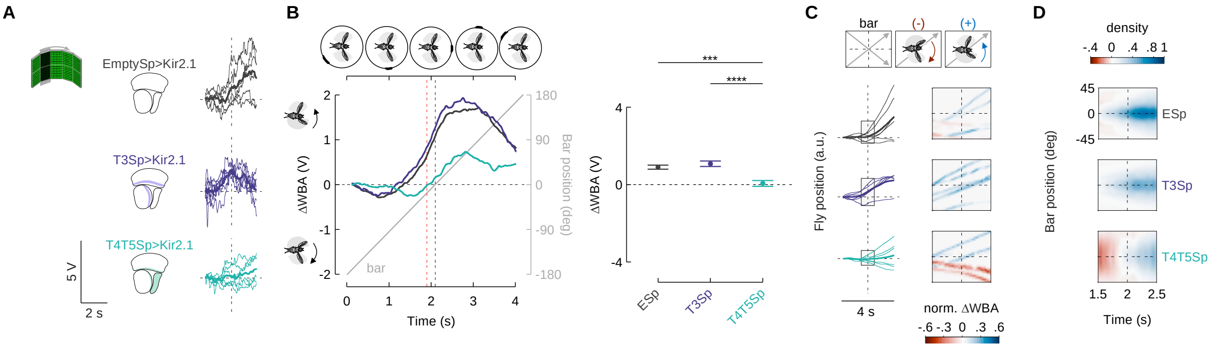


**Supplementary Figure 3.** Hyperpolarization of T3 does not compromise the syn-directional response. (**A**) Left: representation of the visual stimulus (dark bar revolving at 90° $s^{-1}$). Middle: schematic representation of the optic lobe regions where Kir2.1 channels were expressed in the three genotypes tested. Right: single trials (3 repetitions x 2 directions) of ΔWBA responses (thin lines) to rotation of a luminance-defined bar in three representative flies (T4/T5Sp>Kir2.1 data are reproduced from Keleş et al., 2018). Thick line represents the mean (responses to CCW rotations were reflected and pooled with CW responses). Vertical dashed lines indicate when the bar is at the fly's visual midline while the horizontal ones represent ΔWBA = 0. (**B**) Left: population average time series steering responses (mean ± s.e.m.) in the three genotypes tested (T4/T5Sp data replotted from Keleş et al., 2018) to a luminance-defined bar. Gray shaded region (between vertical red and black dashed lines) represents a 200 ms time window before the bar crosses the fly's visual midline (n = 44 EmptySp>Kir2.1, n = 26 T3Sp>Kir2.1, n = 22 T4/T5Sp>Kir2.1). T4/T5Sp>Kir2.1 reduces syn-directional anticipatory steering, whereas T3Sp>Kir2.1 shows normal syn-directional steering. Right: Dot plot average ΔWBA values across the 200 ms time window per trial. Dark dots indicate the mean and the horizontal bars indicate s.e.m. ($F_{(2, 89)} = 14.48$, $p < .0001$; EmptySp vs T3Sp: $p = .94$; EmptySp vs T4/T5Sp: $p < .0001$; T3Sp vs T4/T5Sp: $p < .0001$). (**C**) Left: arbitrary fly position (thin lines) resulting from the integration of ΔWBA values over time in three representative flies. Thick lines represent the mean. Right: space-time plot of the normalized fly position within the gray shaded boxes highlighted to the left. Color-code represents the direction of the steering effort (red: counter-directional; blue: syn-directional). (**D**) Heat maps of flies' steering effort at the population level in the three genotypes as a function of the bar position. EmptySp and T3Sp flies show a strong syn-directional response (blue blob) while T4/T5Sp flies show an early counter-directional response and a weak syn-directional response.

### Supplementary Figure 4

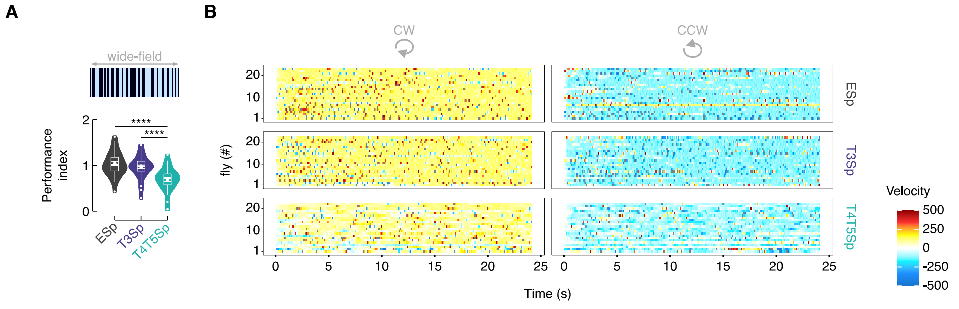


**Supplementary Figure 4.** T3 silencing does not affect the response to the rotation of a wide-field panorama. (**A**) Violin-box plot of the performance index (i.e., gain) to a rotating random wide-field pattern of dark and bright stripes in the three genotypes. T4/T5Sp shows a strong reduction of the smooth tracking gain while T3Sp shows a normal tracking response (pairwise post-hoc comparisons adjusted Bonferroni, EmptySp vs T3Sp: $p = 1$; EmptySp vs T4/T5Sp: $p < .0001$; T3Sp vs T4/T5Sp: $p < .0001$). (**B**) Raster plot of the velocity (yellow-red: CW; green-blue: CCW) per fly during the rotation of the wide-field panorama (each bin represents 100 ms of average velocity). Note that T4/T5Sp>Kir2.1 flies reduce the tracking velocity, whereas T3Sp>Kir2.1 flies show a velocity comparable to control flies (n = 20 EmptySp>Kir2.1, n = 22 T3Sp>Kir2.1, n = 21 T4/T5>Kir2.1).

### Supplementary Figure 5

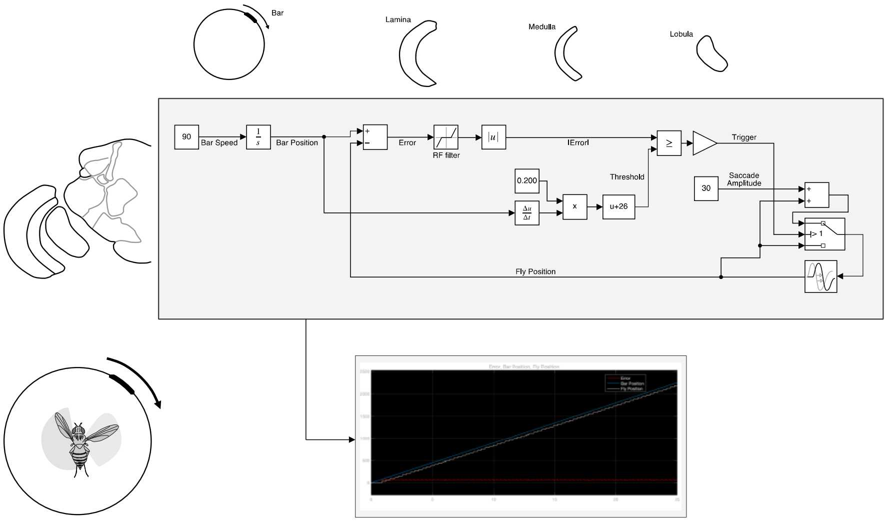


**Supplementary Figure 5.** Control model for triggering saccades. Top: simulink (MATLAB) implementation of the physiologically-inspired model in **Figure 6**. The bar speed (90° $s^{-1}$) is integrated over a quite narrow time window (~ 200 ms). This means that T3 neurons might encode the bar speed and a downstream partner might integrate this information over a selective amount of time, encoding the bar position and triggering a saccade when the amount of calcium reaches a specific threshold. In this model saccade amplitudes are considered a fixed parameter (30°) but in an alternative version they could be easily tuned to the bar speed (as a role played by the T4/T5 pathway). Bottom: simulation of the fly behavior (angular position) in a bar tracking task according to the control model.
